# Consumer resilience suppresses the recovery of overgrazed ecosystems

**DOI:** 10.1101/2025.02.04.636488

**Authors:** Nathan B Spindel, Aaron W. E. Galloway, Julie B. Schram, Gwiisihlgaa Daniel Mcneill, Sgiids Ḵung Vanessa Bellis, Niisii Guujaaw, Jaasaljuus Yakgujanaas, Ondine Pontier, Markus Thompson, Lynn C Lee, Daniel Okamoto

## Abstract

1. Many heterotroph species perish when faced with severe food limitation, others can persist, adapt, and thrive. Sea urchins are emblematic of this paradox: they can overgraze kelp forests to form barren habitats, but can then survive for decades in these nutritionally depauperate seascapes. Understanding the mechanisms enabling persistence under starvation, and rapid recovery when food returns, provides insight into how consumer resilience shapes ecosystem dynamics.
2. We quantified how food abundance, quality, deprivation, and reintroduction influence bioenergetic performance in the red sea urchin (*Mesocentrotus franciscanus*), integrating field observations of kelp forest and barren populations with a controlled feeding experiment. We measured respiration, feeding rates, gonadal growth, and fatty acid biomarkers to test how habitat history and diet jointly govern metabolic plasticity and nutrient assimilation.
3. Resting metabolic rates (RMR) were nearly twofold higher in kelp forest urchins than barrens conspecifics, yet feeding rates were equivalent across habitats, indicating that metabolic depression does not constrain food intake. Reciprocal shifts emerged in the experiment: starvation reduced RMR and lipid reserves in kelp forest urchins, while feeding elevated both traits in barrens urchins to levels comparable with kelp forest conspecifics. These results demonstrate rapid physiological compensation in response to both food deprivation and reintroduction.
4. Diet quality strongly modulated performance. Urchins fed nutritionally poor monospecific diets consumed more biomass and calories than those on diverse, polyunsaturated fatty acid (PUFA)-rich diets, but did so with markedly lower efficiency of conversion to gonadal tissue. Fatty acid assimilation revealed that starvation elevated bacterial and biofilm biomarkers in tissues, whereas algal diets enriched essential PUFA profiles, particularly when diets were diverse. These results highlight that both quantity and quality of food influence consumer recovery trajectories, with nutritional geometry shaping efficiency of energy and nutrient use.
5. Together, our findings show that *M. franciscanus* exhibits pronounced metabolic resilience, allowing persistence in barren habitats and rapid reactivation of grazing and reproduction when food becomes available. This work links nutritional ecology to ecosystem feedbacks by showing how compensatory feeding and metabolic flexibility enable consumers to maintain pressure on primary producers, thereby influencing the stability, hysteresis, and recovery of degraded ecosystems.

## 1. Introduction

Variation in food availability can directly shape the fitness of consumers and associated food web dynamics. Such variation results from, for example, bottom-up changes in primary production (Edwards et al., 2020), shifts in availability of allochthonous food resources (Leal et al., 2022), and/or strong consumer pressure (Gangal et al., 2021). Declines in the quantity or quality of available food can reduce rates of reproduction, juvenile survival, growth, and adult survival in many taxa (Okamoto et al., 2012, Mduma et al., 1999, Olsen et al., 2011). A diverse range of animals across trophic levels, taxonomic groups, and ecosystems (Randall, 1961, Silliman and Zieman, 2001, Mysterud, 2006, Kayal et al., 2012, Ling et al., 2015) are known to periodically graze down and deplete resource productivity leading to barren landscapes of nutritionally poor, scarce, or low productivity food resources. While many fields of study have focused on how these trophic interactions shape population dynamics, less is known about the mechanistic basis of consumer resilience to nutritional deficits. If consumers are to avoid death by starvation, they must emigrate, switch their diet to alternative foods, and/or cope (McCue et al., 2017). Among those that cope, it remains unclear how metabolic plasticity and resilience interact with changes in the abundance and quality of food to shape energetic traits and fitness.

Recent research in nutritional ecology has shown that consumers may regulate intake not simply to maximize calories, but to balance multiple nutrients simultaneously (Simpson et al., 2004, Raubenheimer et al., 2009). Across diverse taxa, from beetles to primates, individuals can adjust their intake of macronutrients to fulfill physiological requirements for individual micronutrients, even at a cost to growth or survival (Jensen et al., 2012, Harrison et al., 2014, Solon-Biet et al., 2014, Tait et al., 2014). Mobile consumer species may respond to reductions in food density or productivity by expanding or shifting their geographic range (Brown and Kotler, 2004) or reducing activity (Storey, 2015). Consumers may also expand their dietary breadth (Tinker et al., 2012) capitalize on exogenous food subsidies (Britton-Simmons et al., 2009), or engage in a last-gasp reproductive bout before dying (Kirkwood and Rose, 1991). In addition to these behavioral shifts, a wide range of eukaryotes avoid minimize the effects of starvation or malnutrition by mobilizing energy reserves for basic functions and/or prioritizing maintenance at the cost of structural growth and/or reproduction (Vilela et al., 2008, Lesage et al., 2001, Chippindale et al., 1993, Ellers, 1995). While heterotrophs can conserve energy and reduce metabolic demands under food deprivation (Storey, 2015, Spindel et al., 2021), the effects of nutritional limitation and resurgence on such metabolic flexibility and the attendant consequences for performance remain poorly studied (McCue et al., 2017). Metabolic rate correlates strongly with specific physiological processes like respiration rate (Glazier, 2015), but it is less clear how metabolic demands feed-back to affect maximum potential food intake and investment in reproduction. For example, while resting metabolic rate (RMR) reflects baseline energetic maintenance, food intake must supply not only energy for homeostasis but also essential nutrients for the growth of reproductive tissues (Sterner and Elser, 2002). A comprehensive understanding of an organism’s metabolic flexibility under food limitation therefore requires integration of energetics with nutrient balance. Beyond the capacity of consumers to modulate metabolic demands under scarcity, the quality and composition of available resources emerge as equally critical determinants of performance and recovery.

Shifts in both resource quantity and quality affect consumer energetics and performance even when caloric intake is sufficient. Macronutrient deficiencies can alter feeding behavior, growth, and longevity in a wide range of organisms (Solon-Biet et al., 2014, Cruz-Rivera and Hay, 2000). Some micronutrients are “essential” in the diets of many consumers (Parrish, 2009). For example, highly unsaturated fatty acids (HUFAs), such as docosahexaenoic acid (DHA), can be critical biomolecules for successful reproduction, growth, and survival (Fuiman and Perez, 2015, Ruiz et al., 2021). Yet many consumers lack the capacity to synthesize these essential nutrients (Twining et al., 2021). When quality of available food is low, consumers may compensate by increasing consumption to meet metabolic and nutritive demands. This phenomenon is a central prediction of nutritional geometry and has been documented across both terrestrial and marine systems (Raubenheimer et al., 2009, Raubenheimer and Simpson, 1993, Cruz-Rivera and Hay, 2000, Berner et al., 2005, Fink and Von Elert, 2006, Jochum et al., 2017). This framework suggests that the balance of nutrients, not energy alone, governs performance and recovery (Raubenheimer and Simpson, 1993, Simpson et al., 2004, Raubenheimer et al., 2009, Simpson and Raubenheimer, 2012).

Sea urchins provide a powerful model to investigate metabolic flexibility under food limitation. They are dominant herbivores that often graze down highly productive and biodiverse algal assemblages (Schiel and Foster, 2015) leaving behind low-productivity barrens which can persist for decades (Filbee-Dexter and Scheibling, 2014). Survival rates of some sea urchins in barrens have been hypothesized to be high as a result of phenotypic plasticity (Edwards and Ebert, 1991). Recent observations demonstrated a 50% decline in RMR between sea urchins occupying food-rich versus food-poor habitats (Spindel et al., 2021). Understanding how such metabolic depression in food-poor habitats affects the capacity of sea urchins to quickly resume consumption and reproduction when food becomes available provides a direct test of theories of metabolic flexibility and nutritional ecology. Moreover, the timescale over which animals respond to resurgent food availability and how different food qualities influence the recovery of reproductive capacity, energy stores, and RMR remains poorly understood (McCue et al., 2017).

We focused on the barren-forming red sea urchin, *Mesocentrotus franciscanus*, because of its important roles in structuring nearshore algal ecosystems, commercial fisheries, and indigenous culture (Rogers-Bennett and Okamoto, 2020). We addressed two objectives: (1) quantifying feeding rates and metabolic responses following starvation as a function of food quality, and (2) characterizing how habitat and diet affect assimilated FA biomarker profiles as a step toward linking experimental nutrition with wild dietary dynamics. Based on large differences in RMR between barren and kelp forest urchins (Spindel et al., 2021), we hypothesized that (a) RMR would be highly flexible as a function of declines and resurgences in food availability, (b) these metabolic shifts would be accompanied by shifts in energy stores and reproductive capacity, and (c) calorically rich but nutritionally poor diets would induce compensatory feeding and yield less efficient recovery than nutritionally-rich diets.

## 2. Materials and Methods

We evaluated the effects of habitat of origin (kelp forest vs. barrens) and experimental diet on both bioenergetics (i.e., metabolism, grazing rate, growth, and gonad production) and FA dietary biomarkers of *M. franciscanus*. Field assays provided natural ecological context for our laboratory experiment and included measurements of RMR, body size specific gonad mass, and FA biomarker profiles in each habitat. We then subjected individuals from neighboring barrens and kelp forest habitats to a six-week aquarium experiment where we controlled their diets and measured bioenergetic responses and FA biomarker profile outcomes.

### Field Collections

We collected individuals using SCUBA from Surge Narrows, British Columbia, (50.22°N, 125.16 °W) from a depth of 2-3 m and 10-12 m relative to mean tide from kelp forest and nearby barrens habitat, respectively (n = 30 per habitat, 60 total; mean test diameter = 69.4 mm ± 1.02 mm SE) in August of 2019. We immediately transported collected individuals in insulated coolers to the Hakai Institute Quadra Island Ecological Observatory (HIQIEO) where we transferred them to flow-through sea tables and allowed them to recover for a period of 48 hours. We collected additional individuals from similar depths for sacrificial sampling to compare habitat-specific gonadal mass allometry and fatty acid composition at Surge Narrows with additional study sites in Gwaii Haanas National Park Reserve, National Marine Conservation Area Reserve, and Haida Heritage Site on Haida Gwaii, including Faraday (52.61° N, 131.49° W) and Murchison (52.60° N, 131.45° W) Islands.

### Aquarium Setup

We used outdoor sea tables supplied with flow-through treated seawater for the laboratory experiment. Seawater was pumped from a depth of 20 m, filtered at 100 µm a. UV sterilized,. We subdivided each sea table into multiple 17L aquaria (n = 60 aquaria) using plastic bins, each supplied with independent flow-through seawater (flow = 2.20 Lhr^-1^; turnover = 3.12 d^-1^) via irrigation plumbing (**figure S1**). Temperature, salinity, and pH were recorded daily (temperature = 13 ± 1.01 SE, salinity = 28.85 ppt ± 0.35 ppt SE, pH = 7.61 ± 0.13 SE). We haphazardly assigned individual urchins to solitary chambers in each aquarium following the recovery period

### Acclimation and Experimental Treatments

We minimized relocation stress by allowing the urchins to acclimate to aquarium conditions for 48 hours prior to initiating treatments. We assigned experimental subjects to one of three diet treatments: 1) Monospecific diet (*Nereocystis luetkeana*), *ad libitum*; 2) Diverse diet (*N. luetkeana, Ulva sp., Chondracanthus corymbiferus,* and *Dilsea californica*), *ad libitum*; and 3) Starvation, resulting in a fully-crossed factorial design (n = 10 urchins per habitat-diet treatment combination). These algal taxa have similar caloric content, C:N ratios, and protein content but substantially different concentrations of nutritionally valuable fatty acids (electronic supplemental material; **table S11**). We collected fresh macroalgae for diet treatments from a depth of 3-6 m at nearshore sites around Quadra Island (*N. luetkeana* at Quathiaski Cove 50.04°N, 125.22 °W, *Ulva sp., C. corymbiferus,* and *D. californica* at Wa Wa Kie Beach 50.04°N, 125.17 °W) and transferred collected algae to flow-through sea tables at the HIQIEO. We prevented decomposition of macroalgal tissue by collecting fresh macroalgae immediately prior to and periodically throughout the experiment, only using algae collected within one week. To standardize composition of fresh macroalgal diets we only used thallus tissue which was free of reproductive patches, edges prone to decomposition, and conspicuous epibionts. We maintained experimental treatments for thirty-three days to allow for controlled diets to be assimilated into new urchin biomass.

### Energetics

#### Respirometry

We quantified respiration rate as a proxy for metabolic rate, focusing on resting metabolic rates (RMR). We selected RMR to avoid the confounding effects of specific dynamic action (Lighton, 2018); to control for this we starved focal subjects for 48 hours prior to respirometry then immediately resumed prescribed feeding after respirometry sessions. We quantified respiration rates using custom-built sealed acrylic chambers (Spindel et al., 2021). To account for the displacement volume of the urchins when calculating respiration rates, we subtracted internal test volume of the urchins (V – calculated from test diameter (D) and height (H) assuming oblate spheroid geometry (V = 4/3π D^2^H)) from the total chamber system volume. We accounted for background oxygen dynamics using an identical empty respiration chamber in the same environmental conditions in parallel with every set of three respiration chambers containing urchins. We conducted reproducible and transparent quality control on oxygen time series data using the R package respR (Harianto et al., 2019).

We standardized RMR with total metabolically active biomass which we measured as ash-free dry mass (AFDM). AFDM represents non-skeletal soft tissue as the difference between dry mass and post-combustion ash mass. We first cracked the test of the urchins and discarded the coelomic fluid, then dried the carcasses for 24 hours at 60 in a drying oven, then weighed the dried carcasses. We measured post-combustion ash mass by combusting dried carcasses for six hours at 450 in a muffle furnace, then weighed the ashes of each carcass (scale resolution =0.01 mg). To contextualize measurements of RMR made on experimental subjects maintained in the laboratory with natural variation, we also sampled the wild population at the beginning and end of the experiment from the same collection site.

#### Feeding Rate and Assimilation Efficiency

We measured food consumption and assimilation efficiency using repeated trials spanning from day 7-14 and 23-33 in shaded sea tables to limit macroalgal growth. For each trial, we measured the wet mass of food at the beginning and endpoints, as well as the mass of feces at the endpoints. We standardized wet weight measurements of algae by spinning thallus tissue in a colander for 30 s prior to recording a mass measurement. We approximated algal dry weights for each of the four species of algae by generating species specific linear regressions between wet and dry weights (electronic supplemental material; **figure S3**). We calculated mean feeding rate as the difference in dry weight of algae from the beginning and end of each feeding cycle divided by the amount of time in the feeding cycle (electronic supplemental material; **method S2).**

#### Gonadal Mass and Food Conversion Efficiency

To quantify shifts in gonadal mass in experimental subjects, we calculated the difference between an estimated initial and measured final state. To estimate initial gonadal mass, we used body size and gonadal mass measurements from representative wild individuals in barrens and kelp forest habitats at Surge Narrows to model the effects of habitat and body size on gonadal dry mass. We measured test volume and gonadal dry mass after heating samples at 60 for 24 h in a drying oven to correct for variation in water content. We sampled a representative size gradient of benthic stage individuals spanning small through large body sizes at Surge Narrows in the barrens and kelp forest habitats at the beginning of the experiment and combined these measurements with identical measurements taken three months prior (Spindel et al., 2021) (kelp forest habitat: n = 49 individuals, minimum test volume = 26 mL, maximum test volume = 668 mL; barrens habitat: n = 59, minimum test volume = 18 mL, maximum test volume = 452 mL).

We quantified food conversion efficiency as a ratio between increase in dry gonadal biomass and total dry food consumption between the beginning and end of the experiment (Brett and Groves, 1979). To infer the beginning habitat specific dry gonadal biomass of experimental subjects for a given body size, we used a model fit to data collected from wild sacrificial samples in the beginning of the experiment collected from the same source populations as the experimental subjects. We used an inverse problem approach to estimate food conversion efficiency, combining error in measurement models estimating treatment specific feeding rates and gonad mass. To estimate treatment specific feeding rates, we fit an error in measurement model to data from replicate feeding trials, combined with an error in measurement model of gonad mass fit to data from wild individuals at the beginning of the experiment and experimental subjects at the end of the experiment.

#### Total Lipid Content

We evaluated changes in biomass-specific energy reserves by comparing the lipid content of gonadal tissue in wild individuals of comparable size to experimental subjects at the beginning of the experiment with the lipid content of gonadal tissue of experimental subjects at the end of the experiment. To prevent degradation of lipids, we immediately flash-froze all harvested tissue samples using liquid nitrogen, then transferred frozen samples to −80 freezers for storage and later lipid extraction and FA composition profiling (see below). To quantify lipid content of gonadal tissue, we extracted lipids from gonadal tissue biopsies (Schram et al., 2018): tissue samples were lyophilized for 48 h then homogenized. Following homogenization, lyophilized tissue samples were digested in 2 mL chloroform, then sonicated, vortexed, and centrifuged (3000 rpm for 5 m) twice in a 4:2:1 chloroform:methanol:0.9% NaCl solution. After each of these cycles, the lower chloroform layer containing organic compounds was pipetted out and pooled. Subsamples were collected from this pooled solution for gravimetric analysis of total lipid content (Kainz et al., 2017).

#### Fatty Acid Composition

To quantify shifts in FA composition associated with nutritional history and diet, we compared the FA composition of lipids extracted from tissue biopsies collected from individuals in their natural habitat and experimental subjects fed controlled diets of known FA composition. We isolated FAs from the lipid solution extracted from tissue samples. To generate a FA-based dietary resource library, we extracted and characterized FAs from thallus tissue samples harvested from the four species of algae supplied in our diet treatments, avoiding epibiota from algal thallus tissue prior to storage. To quantify shifts in FA composition of gonadal tissues, we estimated the state of gonadal tissue FA composition at the beginning of the experiment using gonadal tissue biopsies from wild individuals of comparable size to experimental subjects collected at the same time and from the same place as the experimental subjects (Foster et al., 2015) and compared this estimated initial state against biopsies of gonadal tissue collected from dissected experimental subjects at the end of the experiment.

To quantify the FA composition of tissue samples, we isolated and analyzed FAs from extracted lipids. Briefly, total lipids were extracted using a modified Folch method and derivatized into fatty acid methyl esters (FAME) using methods described in (Schram et al., 2018) and Thomas et al. (2020). Samples were added to chloroform:methanol, sonicated, vortexed, and centrifuged to separate the organic phase, which was removed and evaporated to dryness under nitrogen gas. The extraction process was repeated after resuspending dried lipid in 2 mL of chloroform. To produce FAME, the dried lipid extracts were resuspended in a 1:2 ratio of toluene and 1% sulfuric acid in methanol and incubated at 90°C for 90 minutes. Following transesterification, the samples were allowed to cool, and the acid was neutralized and hexane was added and the samples were vortexed and centrifuged to capture the FAME. Dried FAME were resuspended in 1.5 mL hexane and stored at −20°C. FAME were analyzed with a gas chromatograph-mass spectrometer (Shimadzu GCMS model QP-2020) fitted with a DB-23 column (30 × 0.25 mm × 0.15 μm, Agilent, Santa Clara, CA, USA), using helium as the carrier gas. The heating program described in Taipale et al. (2016) was utilized to ensure sufficient separation between chromatogram peaks. Fatty acids were identified using external standards (GLC 566C, Nu-Chek Prep, Elysian, MN, USA) and mass spectrometry.

#### Replication Statement

In this study, our inferences are drawn at the individual urchin level, as we tested how habitat of origin and diet treatments influence physiological performance and nutritional assimilation in *M. franciscanus*. Treatments were applied at two main scales: (i) habitat of origin (urchins collected from kelp forest vs. barren habitats) and (ii) dietary treatments (starvation, monospecific diet, or diverse algal diet) applied to individual urchins housed in independent flow-through chambers. Replication was based on multiple individuals per treatment, while repeated measurements (feeding rate, assimilation efficiency, fatty acid composition) were taken from the same individuals to improve precision but not as independent replicates. Chamber identity was modeled as a group-level effect to account for potential non-independence.

**Table 1:** Replication schema for laboratory and field-based analyses.

| Scale of inference | Scale at which the factor of interest is applied | Number of replicates at the appropriate scale |
| --- | --- | --- |
| Individual urchins | Habitat of origin (kelp forest vs. barren) | 30 kelp forest urchins, 30 barren urchins in manipulative experiment. An additional 24 individuals from kelp forests and 13 individuals from barrens were sampled at the beginning of the experiment, but not subjected to experimental treatments (total n = 97). |
| Individual urchins | Diet treatments (applied to individuals in independent chambers: starvation, monospecific diet, diverse algal diet) | 10 individuals per diet x habitat combination (total n = 60). An additional 24 individuals from kelp forests and 13 individuals from barrens were sampled at the beginning of the experiment, but not subjected to experimental treatments. |
| Individual urchins | Time (repeated measures: start vs 33 days) | Not independent replicates; subsamples for within-individual precision (total n = 60). |
| Habitat (gonad FA profiling<br>in wild populations) | Depth: 2-3 m and 10-12 m<br>relative to mean tide from<br>kelp forest and nearby<br>barrens habitat, respectively | 3 islands, 2 depths each (2-3<br>m = kelp forest, 10-12 m =<br>barrens), 10-15 urchins per<br>combination (total n = 76). |

### Data Analyses

#### Univariate

We estimated the population level effects of body size, habitat of origin, and diet on bioenergetic and FA outcomes using Bayesian generalized linear models. RMR models also included a group-level effect of respiration chamber and tube feet FA models included a group-level effect of individual to account for repeated measures. We used a Gamma likelihood and log link function to model these continuous positive outcomes. We accounted for uncertainty in model structures by approximate leave-one-out cross validation (loo) and stacking weights (Vehtari et al., 2016, Yao et al., 2018), a method designed to maximize predictive accuracy. To evaluate goodness of fit for our models, we conducted graphical posterior predictive checks, ensuring that model simulations did not systematically differ from empirical data (Gelman et al., 2013) and estimated Bayesian *R^2^* values(Gelman et al., 2019). We evaluated the sensitivity of parameter estimates and model fits to choices of priors by systematically altering priors and refitting models. Subsequently, we tested hypotheses using Bayes Factors (BF) (Kass and Raftery, 1995) computed from marginal likelihoods using the R package bridgesampling (Gronau et al., 2020). We contrasted estimated marginal means (EMMs) to explore differences between levels of factors while accounting for interactions and adjustments of covariates, expressed in terms of highest posterior density (HPD) intervals the R package emmeans (Lenth et al., 2024). We estimated model posteriors using Stan (Team, 2022) via the R (Team, 2024) package brms (Bürkner, 2017) with 16,000 iterations across four chains, discarding the first half of the iterations per chain as a warm-up, resulting in a posterior sample of 32,000 iterations for each outcome. To ensure convergence, we verified that Rhat (the potential scale-reduction factor) was no higher than 1.01 and the minimum effective sample size (*n_eff_*) was greater than 1,000 (Gelman et al., 2013).

We estimated starting body size and habitat specific gonadal biomass of experimental subjects using a model fit to data collected from sacrificing individuals at the beginning of the experiment from the same source populations as the experimental subjects. To evaluate gonad production during the experiment, we combined this estimated initial state with measurements of per capita food consumption and final gonadal biomass. Because both consumption and gonadal measurements were subject to observational error, we implemented a probabilistic framework that explicitly propagated this uncertainty. In this framework, gonad production was modeled as a function of body size, habitat of origin, and diet, while incorporating error terms for both feeding rate and gonad mass to reflect their respective measurement uncertainty (electronic supplemental material; **method S2**).

#### Multivariate

We visualized multivariate patterns associated with habitat of origin and diet using non-metric multidimensional scaling ordination (NMDS) and tested for significance of predictors using a multivariate analysis of variance. We accounted for the compositionality of fatty acid data using Aitchison dissimilarity, which implements a centered log ratio transformation (Aitchison, 1982). NMDS were fit using the package vegan (Oksanen et al., 2007) in R (Team, 2024). Ordination stress and the number of dimensions are reported within each NMDS plot. We fit correlation vectors for constituent fatty acids, such that vector lengths were scaled by correlation coefficients with respect to NMDS ordination scores for each data point. Fitted fatty acid vector directions indicated the ordination space toward which the focal fatty acid changed most rapidly and had the greatest correlation with the ordination configuration (Oksanen et al., 2023). We identified fatty acids that contributed most to the observed multivariate separation by habitat using a SIMPER analysis (Clarke, 2006). To assess significance of multivariate additive and interaction effects of habitat of origin and diet while accounting for heteroskedasticity and unequal sample sizes, we ran W^*^ (Hamidi et al., 2019) tests.

## 3. Results

### Energetics

#### Resting metabolic rate

Resting metabolic rate (RMR) scaled positively with body size and was strongly influenced by both habitat of origin and diet (Figure 1). Wild kelp forest urchins had nearly twice the RMR of barrens conspecifics (ratio = 1.978, 95% HPD: 1.613-2.380). Experimental manipulations produced reciprocal shifts: kelp forest urchins depressed their RMR after 33 days of starvation (ratio = 0.560, 95% HPD: 0.404–0.731) to levels similar to barrens individuals, while barrens urchins increased their RMR when fed to levels comparable with kelp forest conspecifics. After 33 days, habitat differences were no longer detectable: kelp vs. barrens comparisons overlapped zero both under starvation (ratio = 1.135, 95% HPD: 0.749-1.525) and feeding (monospecific ratio = 1.058, 95% HPD: 0.691-1.430; diverse ratio = 0.985, 95% HPD: 0.661-1.342). Feeding produced especially strong effects in barrens individuals: RMR increased more than threefold relative to wild (monospecific ratio = 3.182, 95% HPD: 2.355-4.073; diverse ratio = 3.280, 95% HPD: 2.199-4.431). In contrast, kelp forest urchins responded strongly to starvation but not feeding, with depressed RMR under starvation (ratio = 0.560, 95% HPD: 0.404-0.731) and high values under either diet (monospecific ratio = 3.060, 95% HPD: 2.040-4.177). Effects of monospecific versus diverse algal diets were largely indistinguishable (kelp ratio = 0.872, 95% HPD: 0.586-1.179; barrens ratio = 0.937, 95% HPD: 0.610-1.272). Full statistical comparisons are presented in the electronic supplementary material (Table S1, Figure S4).

**Figure 1.**
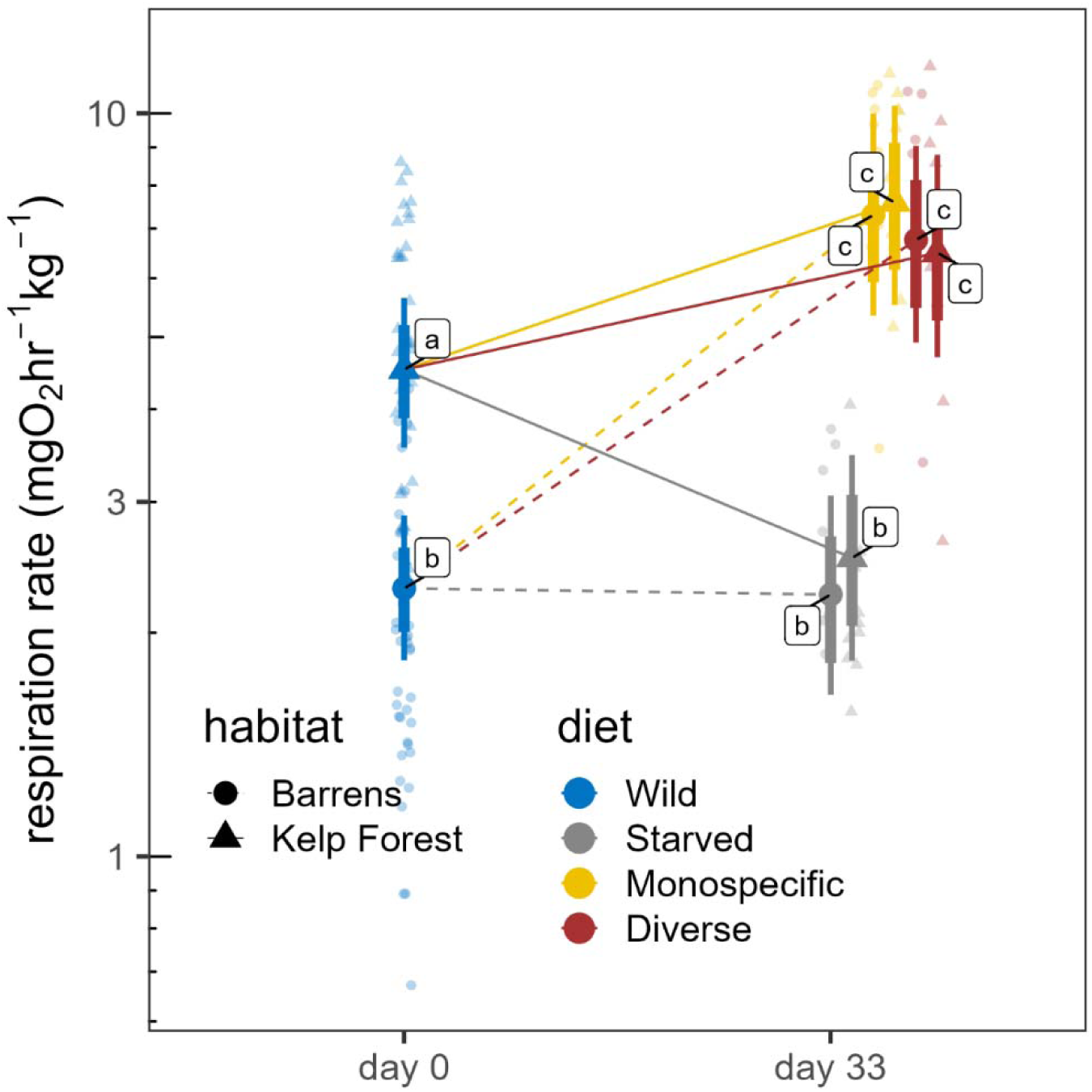
Shifts in resting metabolic rate associated with habitat of origin and diet. Smaller faded symbols represent measured data points, larger solid symbols represent modeled mean values, and thin and thick vertical bars represent 95% and 80% credible intervals, respectively. Lettered labels next to modeled means indicate statistically homogenous groups.

#### Feeding rate

Diet quality was the strongest predictor of consumption (Figure 2). At the start of the experiment, feeding rates did not differ between habitats or diets (ratio = 1.017, 95% HPD: 0.784-1.240). By the end, individuals fed a diverse diet consumed less total biomass and calories than those on a monospecific diet, despite similar per capita intake earlier (ratio = 0.553, 95% HPD: 0.425-0.690). Conversely, polyunsaturated fatty acid (PUFA) intake was initially higher under diverse diets but converged across treatments after 33 days. Statistical model selection details are provided in Tables S2–S4.

**Figure 2.**
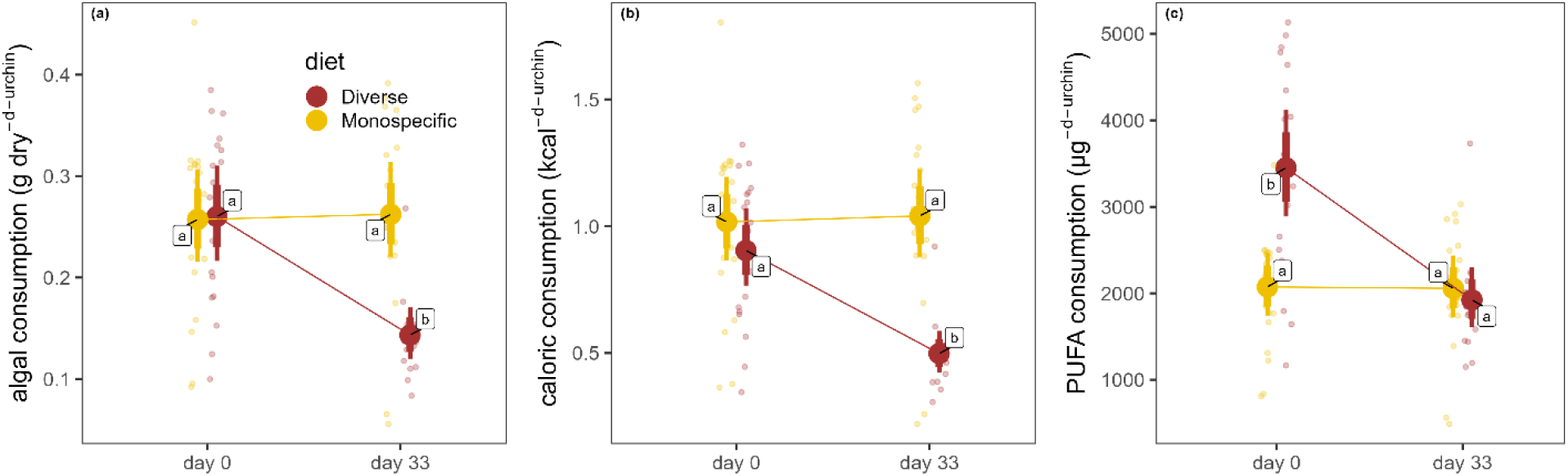
Effects of diet on per capita consumption as a function of time, after controlling for habitat of origin, in terms of (a) algal biomass, (b) calories, and (c) nutritionally valuable polyunsaturated fatty acids (PUFA – fatty acids with two or more double bonds). Smaller faded symbols represent measured data points, larger solid symbols represent modeled mean values, and thin and thick vertical bars represent 95% and 80% credible intervals, respectively. Lettered labels next to modeled means indicate statistically homogenous groups.

#### Assimilation efficiency

Assimilation efficiency was generally stable across treatments. Neither body size, habitat of origin, nor diet produced consistent differences, although there was a slight tendency for efficiency to increase over the course of the experiment (odds ratio = 1.16, 95% HPD 0.86-1.45). These findings suggest that differences in performance between diets were driven primarily by consumption and conversion efficiency, not assimilation itself (Table S5).

#### Gonad mass and lipid content

For a given body size, kelp forest urchins had larger gonads and greater lipid content than barrens conspecifics in the wild (gonad mass: ratio = 1.89, 95% HPD: 1.34–2.46; lipids: ratio = 1.37, 95% HPD: 1.03–1.74). Starvation reduced both metrics regardless of habitat (kelp gonad mass: ratio = 0.66, 95% HPD: 0.40–0.84; kelp lipids: ratio = 0.734, 95% HPD: 0.55–0.93; barrens gonad mass: ratio = 0.66, 95% HPD: 0.45–0.91; barrens lipids: ratio = 0.57, 95% HPD: 0.37–0.77). Feeding restored gonads and lipids in both habitats. In barrens urchins, feeding increased gonadal biomass and lipids relative to wild conspecifics (monospecific: gonad mass ratio = 1.47, 95% HPD: 1.00–1.97, lipids ratio = 1.53, 95% HPD: 1.09–2.01; diverse: gonad mass ratio = 1.38, 95% HPD: 0.95–1.87, lipids ratio = 1.63, 95% HPD: 1.07–2.23). In contrast, fed kelp forest urchins resembled wild conspecifics, suggesting they already had high baseline energy reserves (Figure 3a). Model comparisons supporting these results are reported in Tables S6–S7.

**Figure 3.**
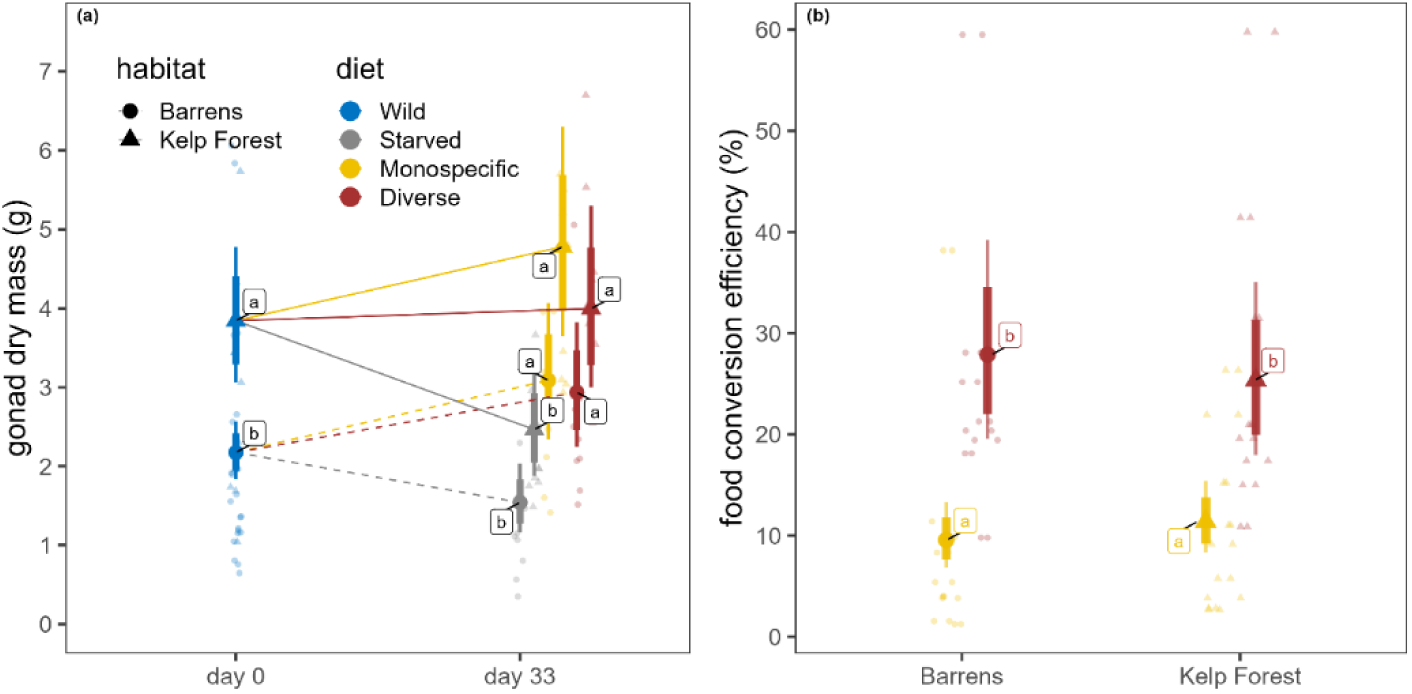
Effects of habitat and diet on (a) gonad mass and (b) food conversion efficiency. Smaller faded symbols represent measured data points, larger solid symbols represent modeled mean values, and thin and thick vertical bars represent 95% and 80% credible intervals, respectively. Lettered labels next to modeled means indicate statistically homogenous groups.

#### Food conversion efficiency

Urchins fed diverse diets converted food to gonadal tissue almost three times more efficiently than those fed monospecific diets, regardless of habitat (ratio = 2.81, 95% HPD: 1.91– 3.81; Figure 3b). This demonstrates that diet quality strongly influences the efficiency of reproductive recovery (Table S8).

#### Fatty acid composition

Urchins from Faraday, Murchison, and Quadra Islands exhibited distinct fatty acid profiles by habitat of origin, and these patterns were sensitive to diet treatments and starvation. Parallel responses were observed in both gonads (figure 4 & 5; electronic supplemental material, figure S7) and tube feet (electronic supplemental material, figure S8). SIMPER analysis identified specific fatty acids associated with habitat and diet (figure 4). Among these, kelp biomarkers (oleic, linoleic, stearidonic, and α-linolenic acids) and biofilm biomarkers (odd-chain fatty acids and the diatom marker palmitoleic acid) were particularly diagnostic. In the field, kelp forest urchins consistently had higher kelp biomarker and lower biofilm biomarker concentrations than neighboring barrens urchins. In the laboratory, controlled macroalgal diets increased kelp biomarker concentrations across habitats. Biofilm biomarkers were similar between starved urchins and barrens urchins in the field, but for kelp forest urchins, starvation more than doubled biofilm biomarker concentrations (ratio = 2.50, 95% HPD = 1.66–3.37).

**Figure 4.**
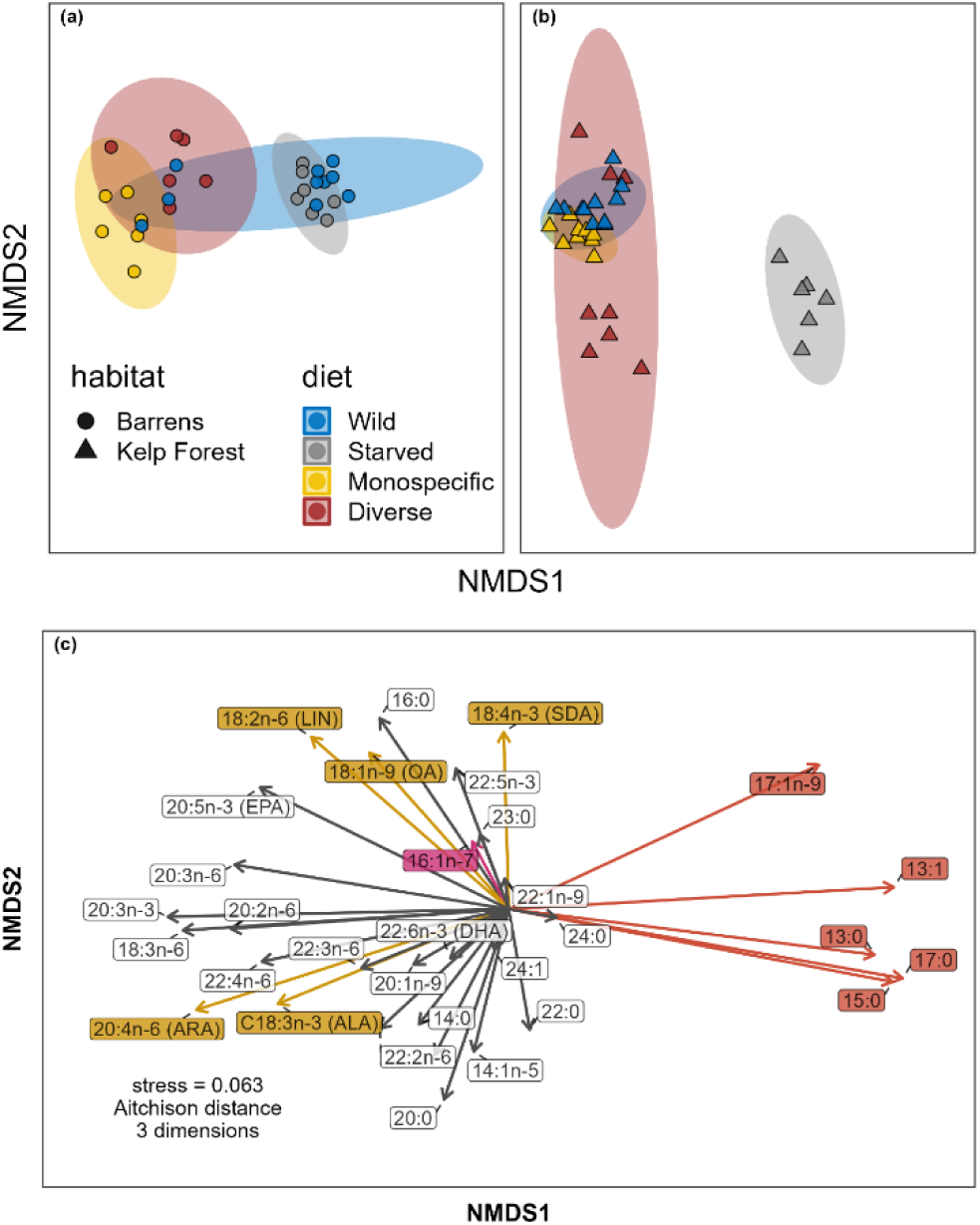
Shifts in gonadal fatty acid (FA) composition as a function of habitat of origin and diet. (a & b) Icons represent individual profiles and ellipses indicate 95% confidence intervals. (c) Arrows indicate correlation vectors for constituent fatty acids, with FAs representing kelp biomarkers, biofilm biomarkers, and diatoms colored in gold, red, and aquamarine, respectively.

**Figure 5.**
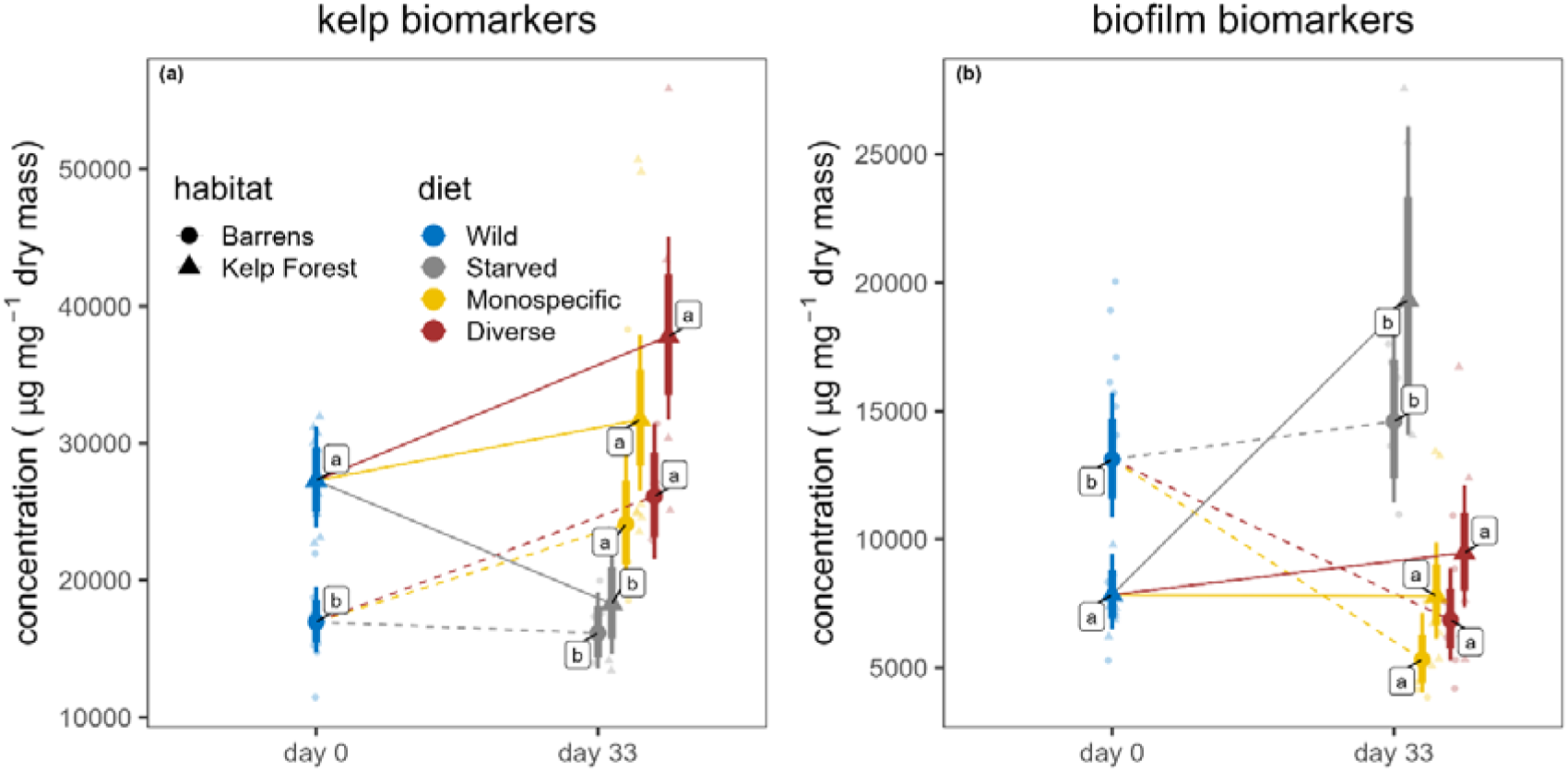
Effects of habitat of origin and diet on gonadal concentrations of (*a*) kelp biomarkers, and (*b*) biofilm biomarkers. Smaller faded symbols represent measured data points, larger solid symbols represent modeled mean values, and thin and thick vertical bars represent 95% and 80% credible intervals, respectively. Lettered labels next to modeled means indicate statistically homogenous groups.

In gonad tissue, fatty acid profiles differed significantly by habitat (W*d = 11.18, p = 0.001, nrep = 999), diet (W*_d = 10.70, p = 0.001, nrep = 999), and their interaction (W*d = 18.00, p = 0.001, nrep = 999). Kelp biomarker concentrations were 27.5% higher in urchins fed a monospecific diet (ratio = 1.28, 95% HPD = 1.07–1.48) and 45.5% higher in those fed a diverse diet (ratio = 1.46, 95% HPD = 1.22–1.70) relative to wild individuals. For biofilm biomarkers, credible intervals for differences between barrens urchins in the field and experimentally starved urchins overlapped zero. Additional details of fatty acid assimilation and trophic modification are provided in the electronic supplemental material.

## Discussion

Our study demonstrates that *M. franciscanus* exhibits striking physiological plasticity in response to variation in food quantity and quality. RMR depressed under starvation but rebounded rapidly when food became available, while feeding rates and tissue composition were strongly shaped by diet quality. Individuals fed diverse algal diets converted food to gonadal tissue more efficiently than those on monospecific diets, and FA profiles confirmed assimilation of complementary nutrients. Together, these results provide empirical evidence for principles long emphasized in nutritional ecology: consumers respond not only to the quantity of food but also to its nutrient balance (Raubenheimer and Simpson, 1993, Simpson et al., 2004, Raubenheimer et al., 2023).

Classic models of consumer resource dynamics predict that overgrazing leads to consumer population decline, potentially allowing resources to recover. However, the ability of *M. franciscanus* to enter hypometabolic states and then rapidly restore RMR and gonadal mass when food reappears suggests an alternative: consumers can persist in food-poor environments and resume grazing with little delay once resources return. This echoes prior work showing metabolic depression and recovery in sea urchins (Dolinar and Edwards, 2021, Okamoto, 2014) and underscores the role of compensatory physiological responses in stabilizing consumer populations.

Compensatory feeding emerged as a central mechanism in our study. Urchins maintained similar caloric intake across treatments but required higher consumption on monospecific, lower-quality diets to achieve reproductive recovery. This pattern parallels findings across diverse taxa where consumers regulate intake of multiple nutrients, not just energy (Simpson and Raubenheimer, 2012, Ruiz et al., 2021, Jensen et al., 2012). The efficiency gains observed in urchins fed diverse diets are consistent with nutritional geometry theory, which predicts that complementary nutrients reduce the need for compensatory overconsumption (Simpson et al., 2004, Raubenheimer and Simpson, 2018). In this way, diet quality mediates the balance between energetic costs and reproductive performance.

Our FA analyses reinforce these conclusions by linking metabolic performance to assimilated nutrient profiles. Wild kelp forest urchins had higher concentrations of kelp-derived FA biomarkers, while barrens and starved individuals showed enrichment of bacterial and biofilm markers. Feeding on diverse macroalgal diets shifted FA profiles toward kelp biomarkers, confirming direct assimilation of high-value nutrients. These results extend prior work tracing FA assimilation in echinoids (Schram et al., 2018, Raymond et al., 2014) and link starvation to elevated microbial biomarkers. Our FA data show that experimentally starved kelp forest urchins and wild barrens urchins elevate bacterial/biofilm biomarkers (odd-chain and branched FAs; palmitoleic acid 16:1n-7) relative to kelp forest conspecifics, consistent with a greater reliance on microbial substrates in food-poor habitats. This finding aligns with prior FA-based reports suggesting the occurrence of benthic diatoms in the diets of urchins found in barrens (Kelly et al., 2008). Parallel microbiome studies show that urchin gut communities shift with habitat; notably, some taxa abundant in *M. franciscanus* guts, such as *Achromobacter*, are rare or absent in their available foods (Miller et al., 2021). This genus can produce protein-rich biomass, illustrating a plausible route by which microbial associates contribute nitrogen- and amino acid-rich nutrition when macrophyte foods are limiting (Ehsani et al., 2019). In addition, *Campylobacteraceae* dominate the guts of *M. franciscanus*, and predictive metagenomics suggest these bacteria metabolize lipids, amino acids, and carbohydrates, potentially enhancing host persistence in food-poor habitats (Hakim et al., 2019, Hakim et al., 2016). Classic experiments corroborate nutritional provisioning by gut microbes: suppressing gut bacteria reduced incorporation of essential amino acids into gonads (Fong and Mann, 1980), and nitrogen-fixing *Vibrio spp.* have been isolated from urchin guts (Guerinot and Patriquin, 1981). Finally, the presence of odd- and branched-chain FAs in urchin tissues, particularly in detritus-feeding contexts, has long been attributed to microbial sources, reinforcing our interpretation of FA shifts under starvation and in barrens as signals of microbial and/or detrital reliance (Zhukova, 2023, Hayashi and Takagi, 1977).

If consumers are able to depress energetic demands in the absence of food, but rebound when food becomes abundant, such phenomena are likely to inhibit such cyclic dynamics and facilitate persistent barren states. Our results demonstrate metabolic resilience of the dominant nearshore grazer, *M. franciscanus* with strong consequences for grazing effects in nearshore ecosystems. While starvation induces metabolic depression, individuals presented with food can quickly recover to feed at consistent rates regardless of whether it has a history of metabolic depression (**figures 1 and 2**). In this study, emaciated individuals were capable of nearly complete recovery of gonadal mass in just over one month when supplied ample food. Moreover, recovery was more efficient with a diverse diet (**figure 3**), supporting the hypothesis that consumers balance the intake of complementary nutrients (Raubenheimer et al., 2023) rather than selecting singular substitutable food resources based on energy density (Houston and McNamara, 2014).

The persistence of barren habitats has often been attributed to grazer abundance (Ling et al., 2015, Filbee-Dexter and Scheibling, 2014), yet our results suggest that grazer nutritional ecology also contributes. Metabolic depression prolongs survival during scarcity, while compensatory feeding on low-quality foods can maintain energy balance once early colonizing algae or microbial biofilms appear. This dynamic creates hysteresis wherein nutritionally poor but abundant foods can sustain populations at high density, suppressing recovery of more diverse, nutrient-rich macroalgal assemblages. Comparable dynamics have been documented in other systems where food quality determines consumer efficiency and population impacts (Marzetz et al., 2017, Foster et al., 2015). Our findings therefore broaden the mechanistic basis for barren stability from consumer density alone to include nutrient-specific feedbacks in grazer-producer interactions.

Although our study was framed in nutritional ecology, the findings have implications for management. The remarkable resilience of *M. franciscanus* suggests that barren formation reduces the resilience of kelp forests by maintaining high consumer capacity to exploit resurgent algae. This aligns with observations that urchin overgrazing exacerbates climate-driven kelp declines (Smale, 2020, Ling et al., 2015). Restoration strategies such as predator recovery (Watson and Estes, 2011, Burt et al., 2018), targeted urchin removals (Miller et al., 2022), and/or commercial urchin ranching (Powell et al., 2020, Angwin et al., 2022) may therefore be necessary to reset the system by reducing consumer density and competition. Importantly, our results suggest that interventions which shift the nutritional landscape (e.g., enhancing the diversity of macroalgal recruitment) could also reduce compensatory feeding pressure and aid kelp recovery.

## Ethics

This work did not require ethical approval from a human subject or animal welfare committee.

## Data, code, and materials

Datasets, code, and electronic supplementary material will be made available to reviewers upon request, and pending acceptance will be archived on an open-access repository preferred by the journal.

## Declaration of AI use

We have not used AI-assisted technologies in creating this article.

## Supporting information

Electronic Supplemental Material

## Acknowledgements

We thank Markus Thompson of Thalassia for assistance with specimen collection and underwater photography at Quadra Island, Brenna Collicutt of the Hakai Institute for technical support; Eric Peterson and the Hakai Institute for providing laboratory facilities at Quadra Island and for technical support; Reyn Yoshioka for assistance with fatty acid analysis at the Oregon Institute of Marine Biology.

