## Supplementary material for "Consumer resilience suppresses the recovery of overgrazed ecosystems": Electronic Supplemental Material

Supplemental methods

Method S1. Feeding rate and assimilation efficiency continued). We limited the growth of macroalgae in dietary treatments by shading the sea tables throughout the experiment using shade cloth. Additionally, we measured growth of each species of algae in the diet treatments in chambers without urchins in parallel with feeding chambers and corrected calculated feeding rates by subtracting algal growth from the post-cycle algal dry weight measurement. We measured feces by stirring tank water to make a homogenous mixture then siphoning the resulting slurry onto a 400 µm sieve. We transferred fecal pellets to pre-weighed and dried glass fiber filter disks, then vacuum filtered for one minute while irrigating with deionized water to remove salt. We then dried the loaded disks at 60 ℃ for 24 h in a drying oven and calculated the difference between the loaded and original disk dry weight. Using the fecal mass and estimated ingested mass we calculated assimilation efficiency as the percentage of dry mass egested relative to ingested during the cycle (49).

Method S2. Inverse problem: modeling gonad mass and feeding rates jointly

1. Gonad mass model:

Equation S1

$$g_{i}^{*}= \beta_{0}^{g}+ \beta_{1}^{g}\log{body size}_{i}+\beta_{2}^{g}H_{i}+\beta_{3}^{g}D_{i}+\beta_{4}^{g}\left( H_{i}\times D_{i} \right)+\beta_{5}^{g}f_{i}^{*}+\epsilon_{i}^{g}$$

Where:

- $g_{i}^{*}$ is the true (latent) gonad mass for the $i$th individual.
- $\log({body size}_{i})$ is the log-transformed body size for the $i$th individual.
- $H_{i}$ and $D_{i}$ are the categorical variables for habitat and diet, respectively.
- $f_{i}^{*}$ is the true (latent) feeding rate for the $i$th individual.
- $\epsilon_{i}^{g}$ is the residual error term for the gonad mass model.
- $\beta_{0}^{g}$ is the intercept for the gonad mass model, and $\beta_{1}^{g},\beta_{2}^{g},\beta_{3}^{g},\beta_{4}^{g},\beta_{5}^{g}$ are the regression coefficients for the corresponding predictors.

The observed gonad mass $g_{i}$ is related to the latent gonad mass $g_{i}^{*}$ by:

Equation S2

$$g_{i}= g_{i}^{*}+{measurement error}_{i}^{g}$$

2. Feeding rate model:

Equation S3

$$f_{i}^{*}= \beta_{0}^{f}+ \beta_{1}^{f}\log{body size}_{i}+\beta_{2}^{f}H_{i}+\beta_{3}^{f}D_{i}+\beta_{4}^{f}\left( H_{i}\times D_{i} \right)+\epsilon_{i}^{f}$$

- $f_{i}^{*}$ is the true (latent) feeding rate for the $i$th individual.
- $\log({body size}_{i})$ is the log-transformed body size for the $i$th individual.
- $H_{i}$ and $D_{i}$ are the categorical variables for habitat and diet, respectively.
- $f_{i}^{*}$ is the true (latent) feeding rate for the $i$th individual.
- $\epsilon_{i}^{f}$ is the residual error term for the feeding rate model.
- $\beta_{0}^{f}$ is the intercept for the feeding rate model, and $\beta_{1}^{f},\beta_{2}^{f},\beta_{3}^{f},\beta_{4}^{f},\beta_{5}^{f}$ are the regression coefficients for the corresponding predictors.
- The observed gonad mass $g_{i}$ is related to the latent gonad mass $g_{i}^{*}$ by:

Equation S4

$$f_{i}= f_{i}^{*}+{measurement error}_{i}^{f}$$

Supplemental figures

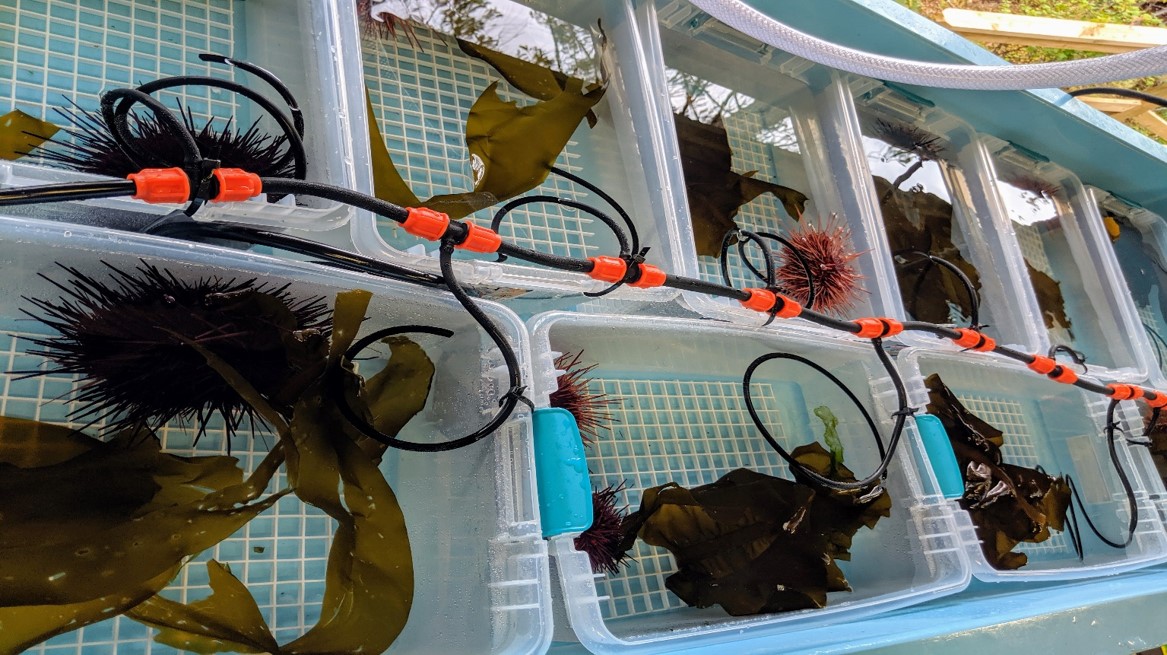

Figure S1: Aquaria array with flow-through seawater supplied to each aquarium. Photo by NBS.

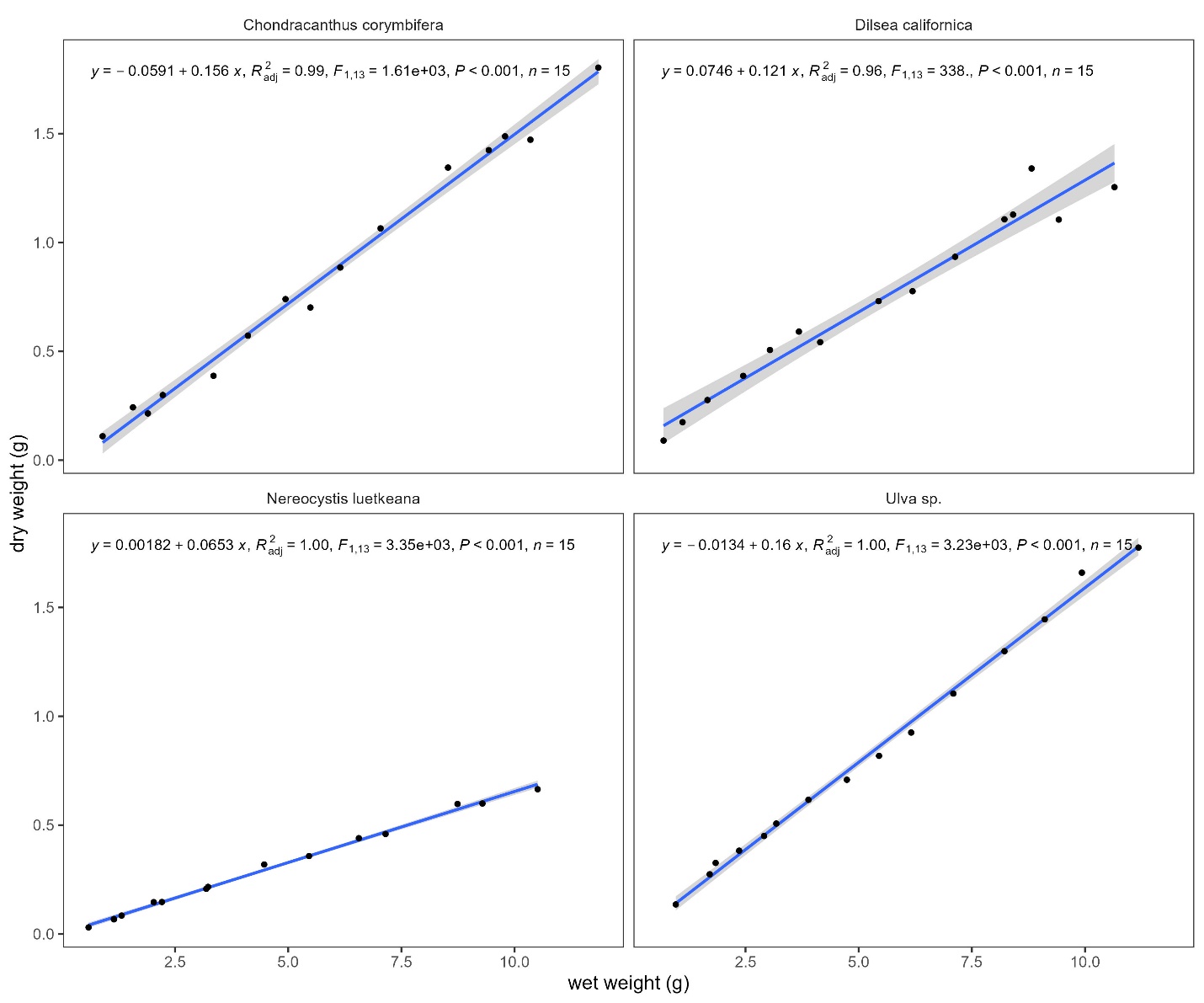

Figure S2. Relationship between wet and dry weight for each algal species supplied in dietary treatments.

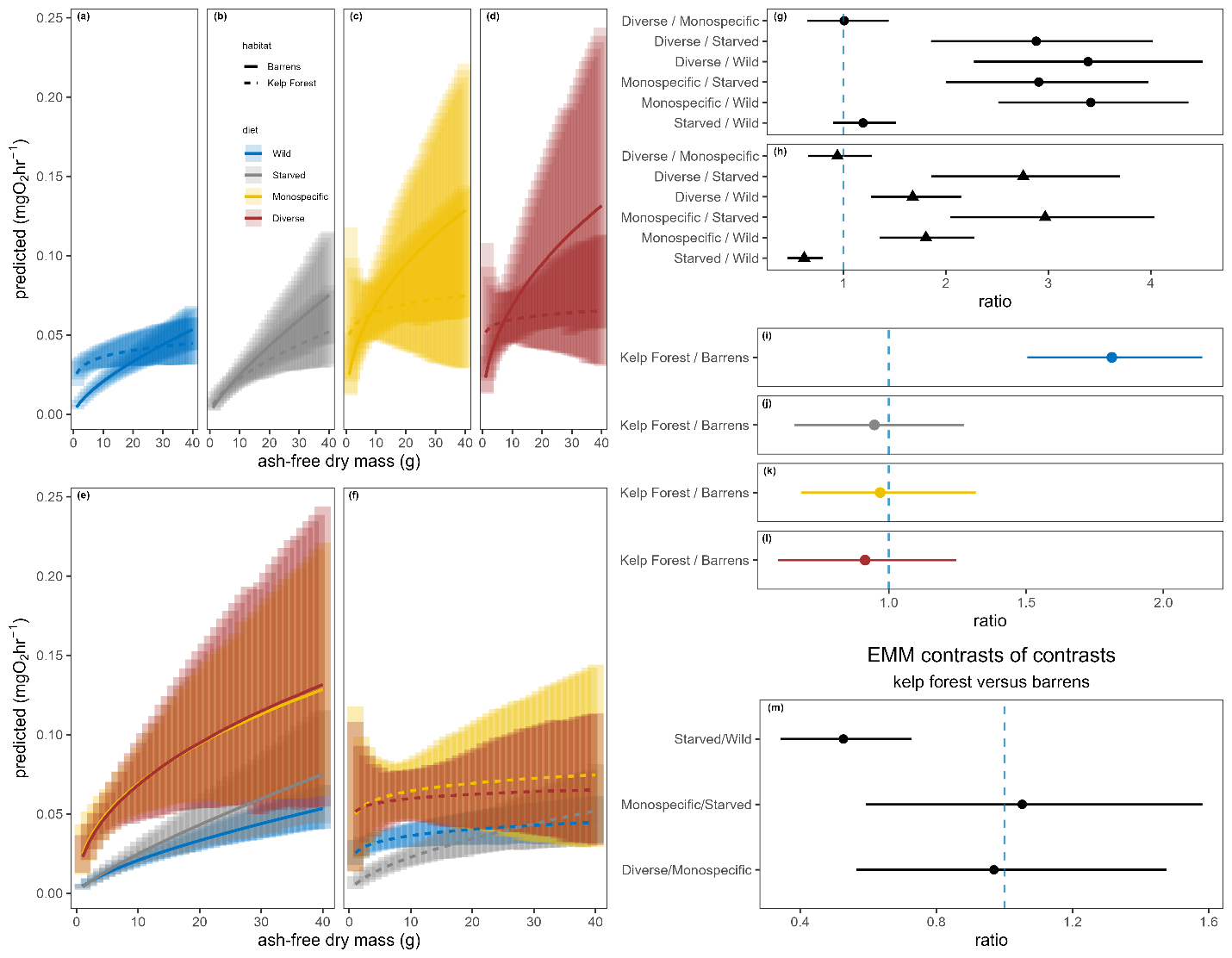

Figure S3. Post hoc respiration analysis using estimated marginal means (EMMs). (a-f) Effects of body size (ash-free dry mass), habitat of origin, and diet on respiration rate. Faded vertical bars represent 95% confidence intervals for predictions at discrete body sizes. (g-m) Contrasts expressed as ratios. Vertical dashed blue line highlights a ratio of one, or no difference. Horizontal bars represent 95% highest probability density intervals. (g-l) Pairwise contrasts by factor. (g) Barrens urchins diet contrasts. (h) Kelp forest urchins diet contrasts. (m) Interaction contrasts (i.e., contrasts of contrasts).
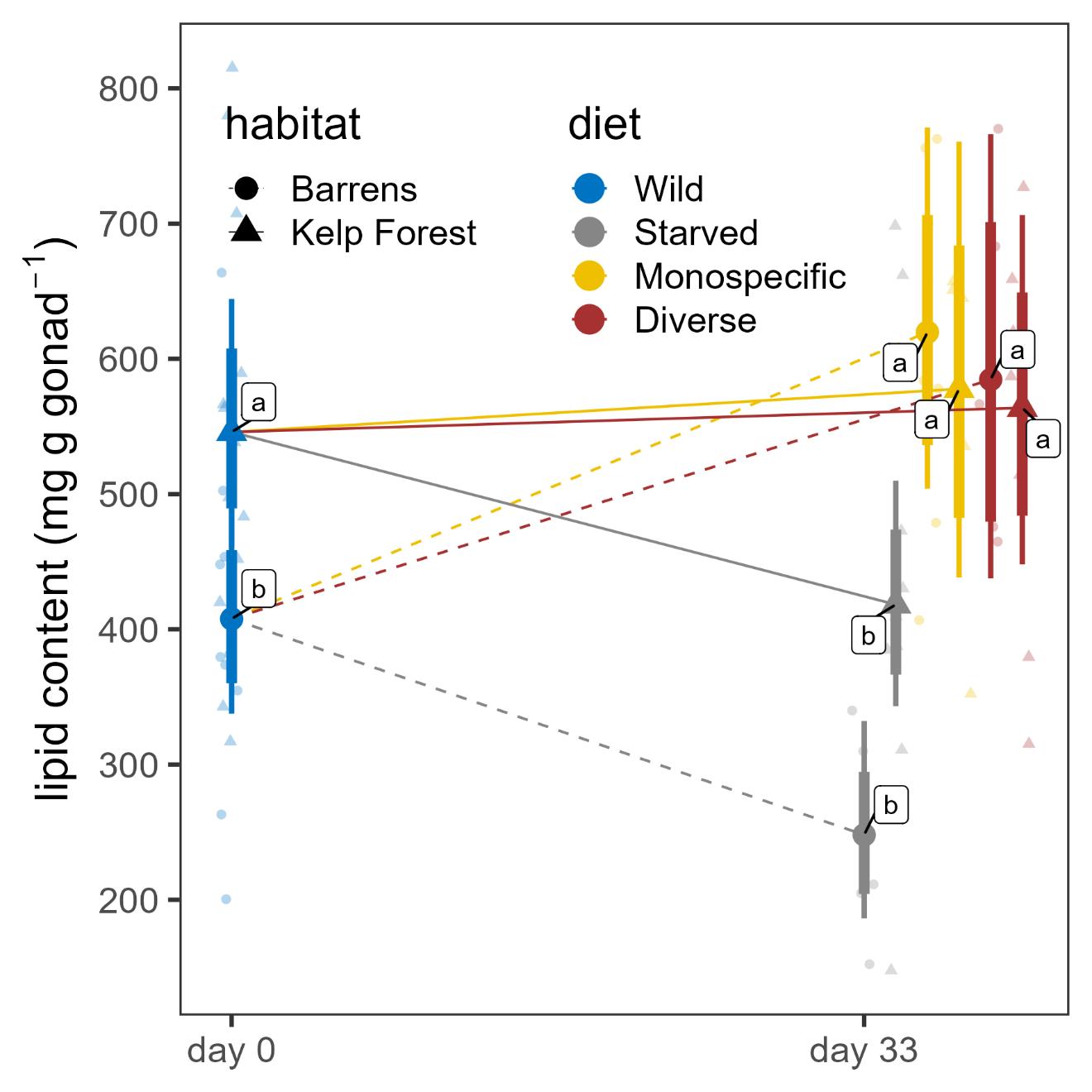

Figure S4. Conditional effects of habitat and diet on gonadal lipid content after controlling for body size (log-test volume). Smaller faded symbols represent measured data points, larger solid symbols represent modeled mean values, and thin and thick vertical bars represent 95% and 80% credible intervals, respectively. Lettered labels next to modeled means indicate statistically homogenous groups.

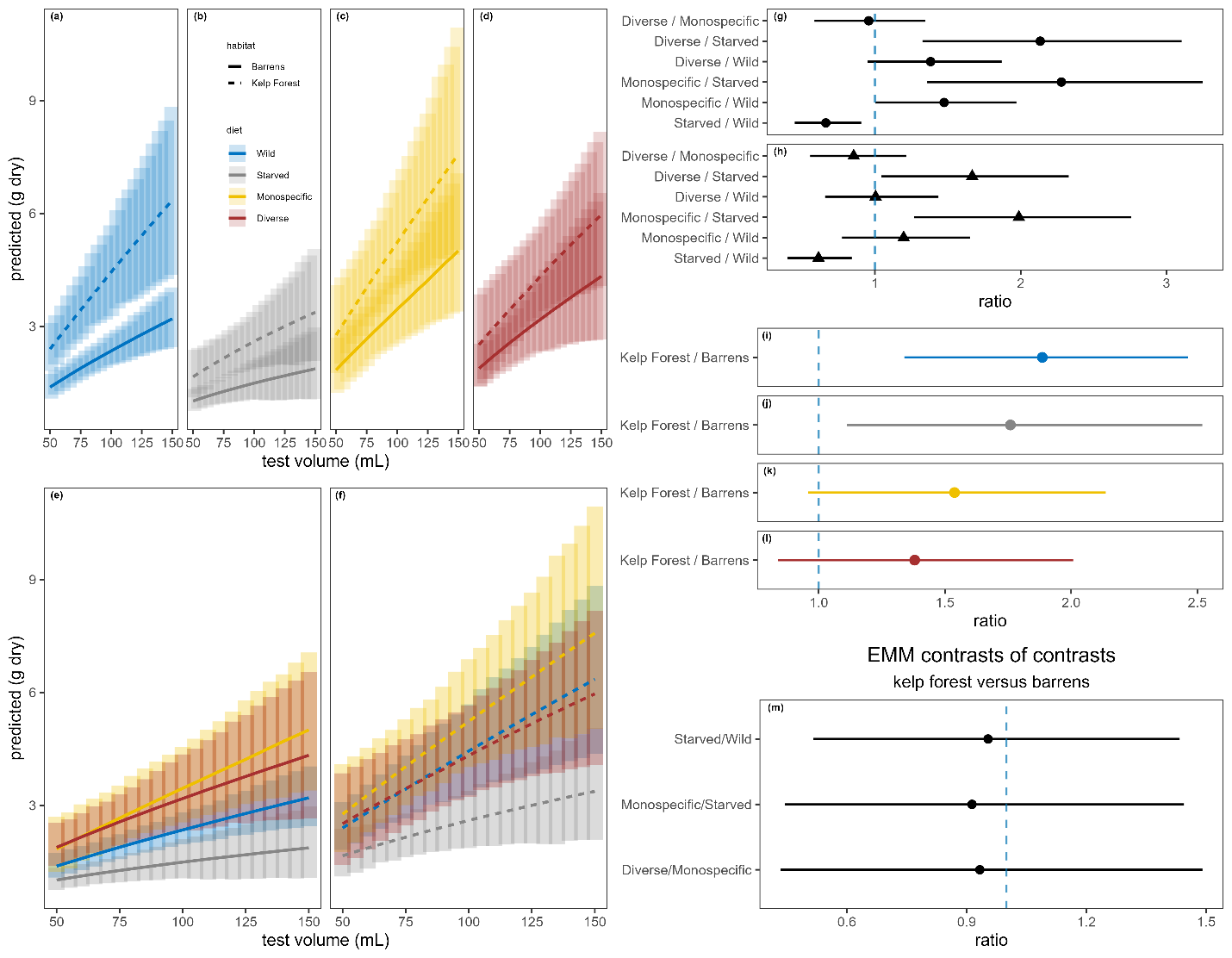

Figure S5. Post hoc gonad mass analysis using estimated marginal means (EMMs). (a-f) Effects of body size (test volume), habitat of origin, and diet on gonad mass. Faded vertical bars represent 95% confidence intervals for predictions at discrete body sizes. (g-m) Contrasts expressed as ratios. Vertical dashed blue line highlights a ratio of one, or no difference. Horizontal bars represent 95% highest probability density intervals. (g-l) Pairwise contrasts by factor. (g) Barrens urchins diet contrasts. (h) Kelp forest urchins diet contrasts. (m) Interaction contrasts (i.e., contrasts of contrasts).

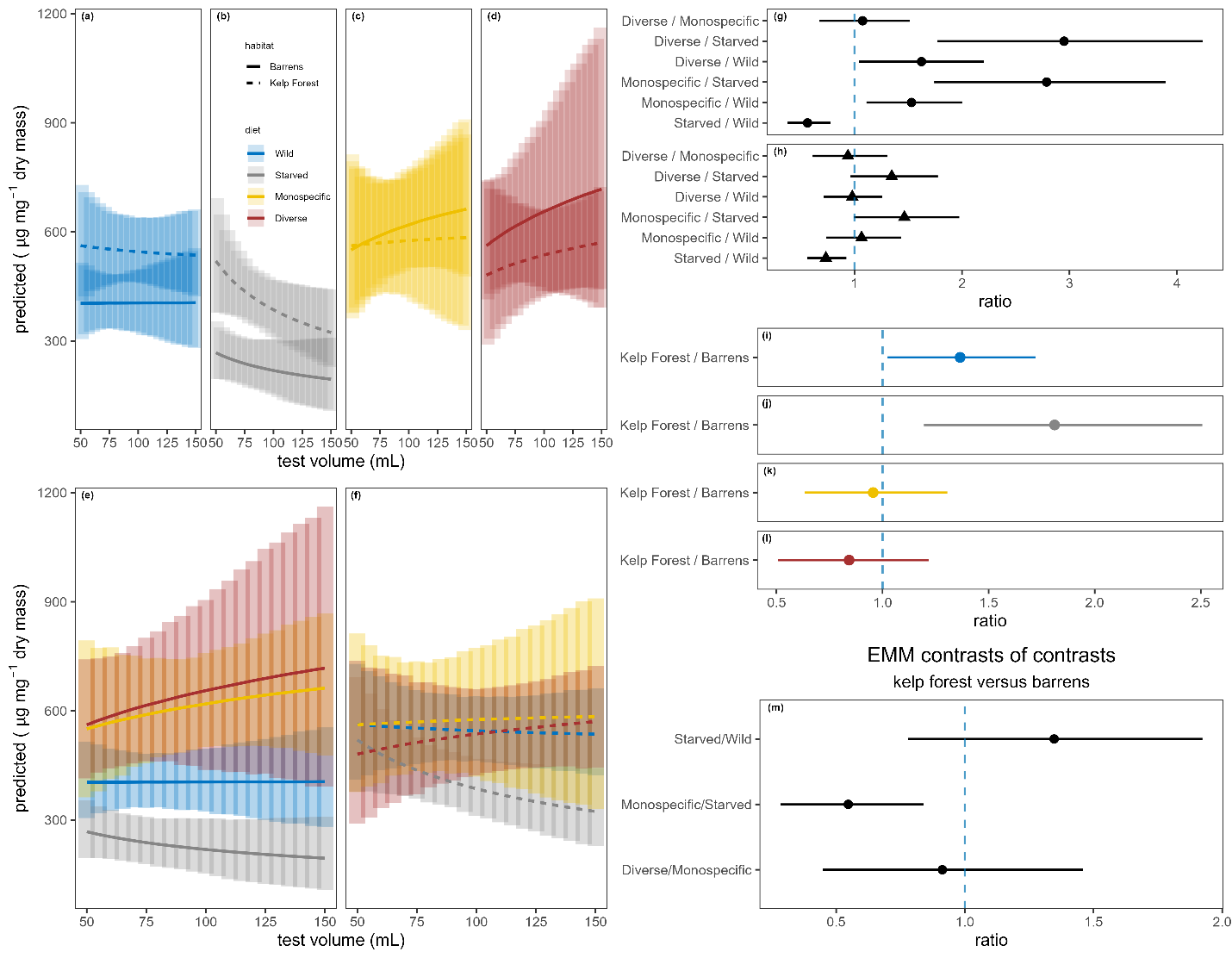
Figure S6. Post hoc lipid analysis using estimated marginal means (EMMs). (a-f) Effects of body size (test volume), habitat of origin, and diet on lipid concentration in gonad tissue. Faded vertical bars represent 95% confidence intervals for predictions at discrete body sizes. (g-m) Contrasts expressed as ratios. Vertical dashed blue line highlights a ratio of one, or no difference. Horizontal bars represent 95% highest probability density intervals. (g-l) Pairwise contrasts by factor. (m) Interaction contrasts (i.e., contrasts of contrasts).

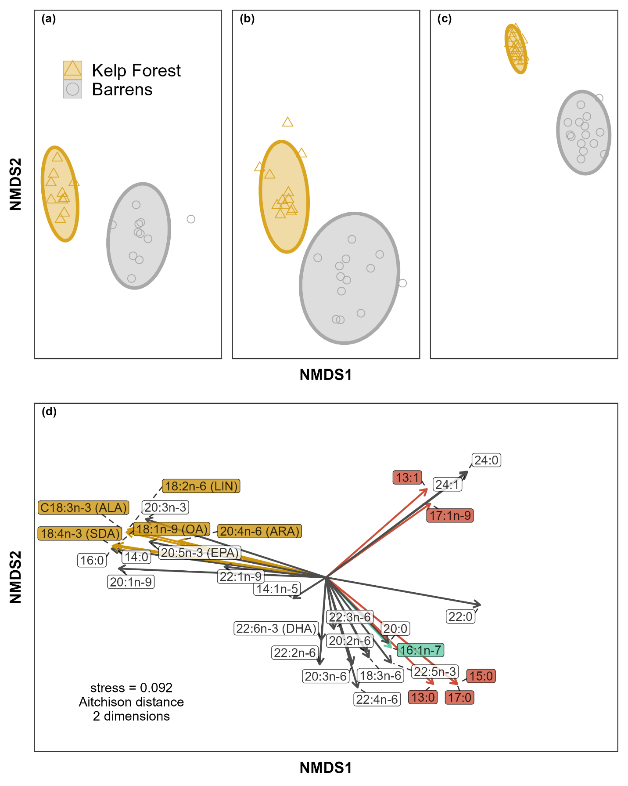

Figure S7. Gonadal fatty acid composition as a function of habitat in wild individuals from three sites in British Columbia, Canada: (a) Surge Narrows (50.22◦ N, 125.16◦ W) between Quadra and Maurelle Islands (here labelled “Quadra”), (b) Faraday (52.61◦ N, 131.49◦ W) and (c) Murchison (52.60◦ N, 131.45◦ W) in Gwaii Haanas on Haida Gwaii. (a-d) Plots illustrate a two-dimensional multivariate ordination of untransformed fatty acid profiles in wild individuals by habitat calculated via non-metric dimensional scaling (NMDS) using Aitchison dissimilarity, which accounts for compositionality in proportional data. (a-c) Faded icons represent individual urchin fatty acid profiles and ellipses indicate 95% confidence intervals. (d) Arrows indicate fitted correlation vectors for constituent fatty acids representing the top 30 contributors to habitat specific differences as indicated by SIMPER analysis, where each arrow’s length is scaled to its correlation with the ordination scores for each data point. Colored fatty acid vectors represent putative biomarkers: gold for kelp biomarkers, magenta for diatom biomarkers, and red for bacterial biomarkers. A total of thirty-four unique fatty acids composed each profile and were used for ordination. The subsets of fatty acids visualized with vectors were selected for their significant contribution to multivariate difference and for their biological relevance to benthic aquatic ecosystems(58).

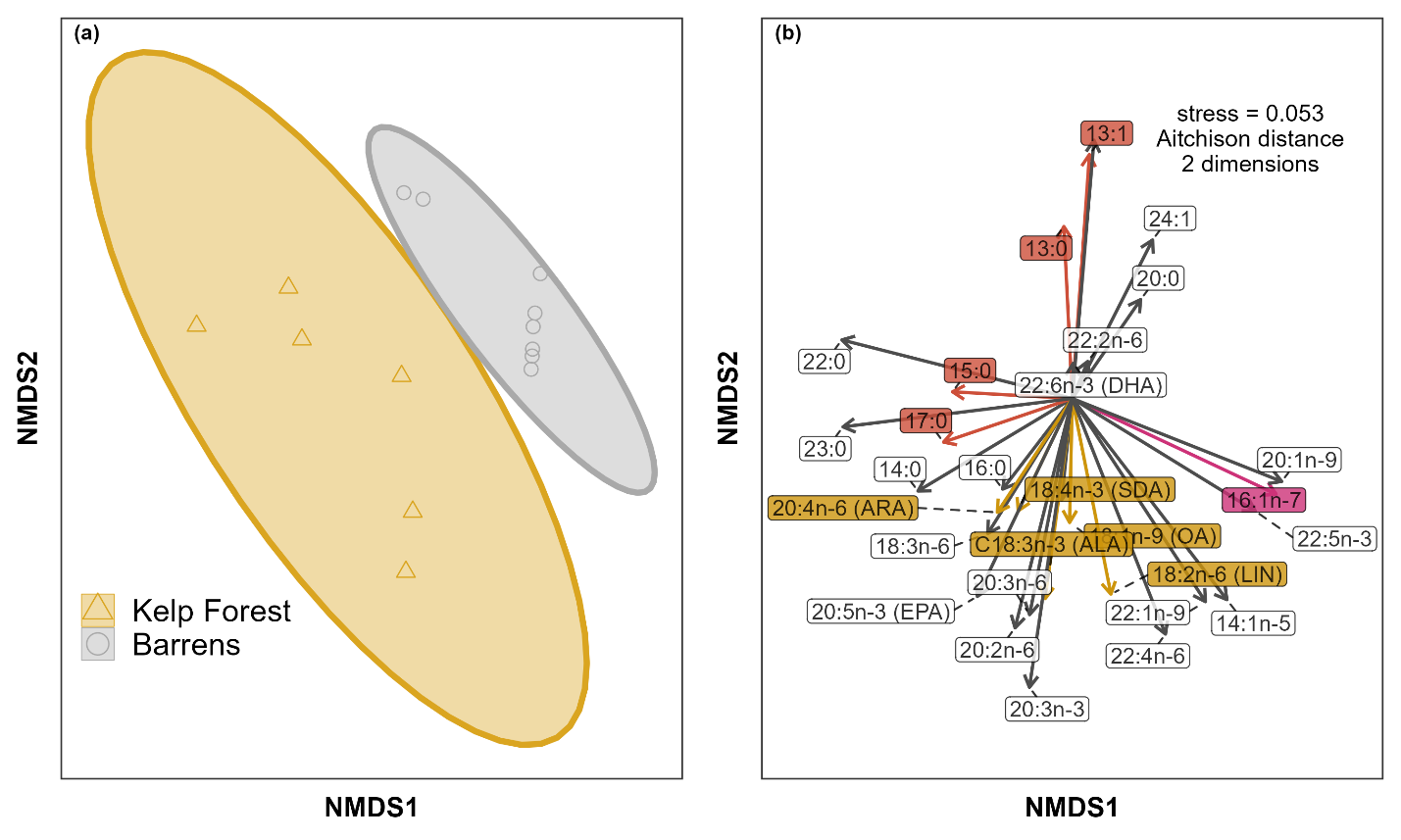

Figure S8. Tube feet fatty acid composition as a function of habitat in wild individuals from Surge Narrows (50.22◦ N, 125.16◦ W) between Quadra and Maurelle Islands. Plots illustrate a two-dimensional multivariate ordination of untransformed fatty acid profiles in wild individuals by habitat calculated via non-metric dimensional scaling (NMDS) using Aitchison dissimilarity, which accounts for compositionality in proportional data. (a) Faded icons represent individual urchin fatty acid profiles and ellipses indicate 95% confidence intervals. Arrows indicate fitted correlation vectors for constituent fatty acids representing the top 30 contributors to habitat specific differences as indicated by SIMPER analysis, where each arrow’s length is scaled to its correlation with the ordination scores for each data point. Colored fatty acid vectors represent putative biomarkers: gold for kelp biomarkers, magenta for diatom biomarkers, and red for bacterial biomarkers. A total of thirty-four unique fatty acids composed each profile and were used for ordination. The subsets of fatty acids visualized with vectors were selected for their significant contribution to multivariate difference and for their biological relevance to benthic aquatic ecosystems(1).

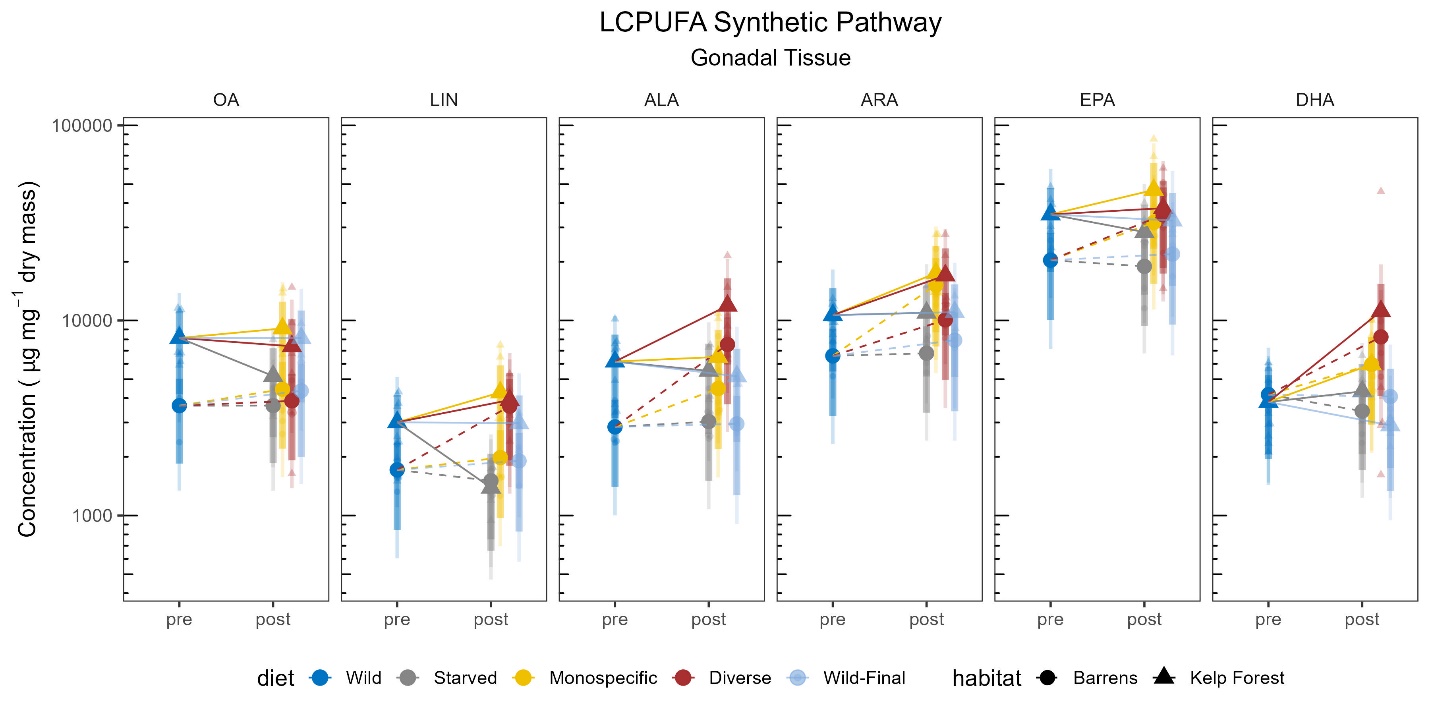

Figure S9: LCPUFA precursor assimilation – gonads. Comparison of the concentration of six focal LCPUFA in gonadal tissue by treatment. Each column of panels represents data for a single focal fatty acid indicated by its abbreviation (left to right: OA, LIN, ALA, ARA, EPA, and DHA). Biosynthesis of these fatty acids generally occurs in sequence from left to right(12). Shapes indicate habitat of origin. Colors indicate diet treatments, where smaller faded symbols represent measured data points, larger solid symbols represent modeled mean values, and thin and thick vertical bars represent 95% and 80% credible intervals, respectively. The horizontal axis is delimited by two time points: “pre”, immediately prior to the experiment (wild individuals only), and “post”, 33 days of treatment exposure.

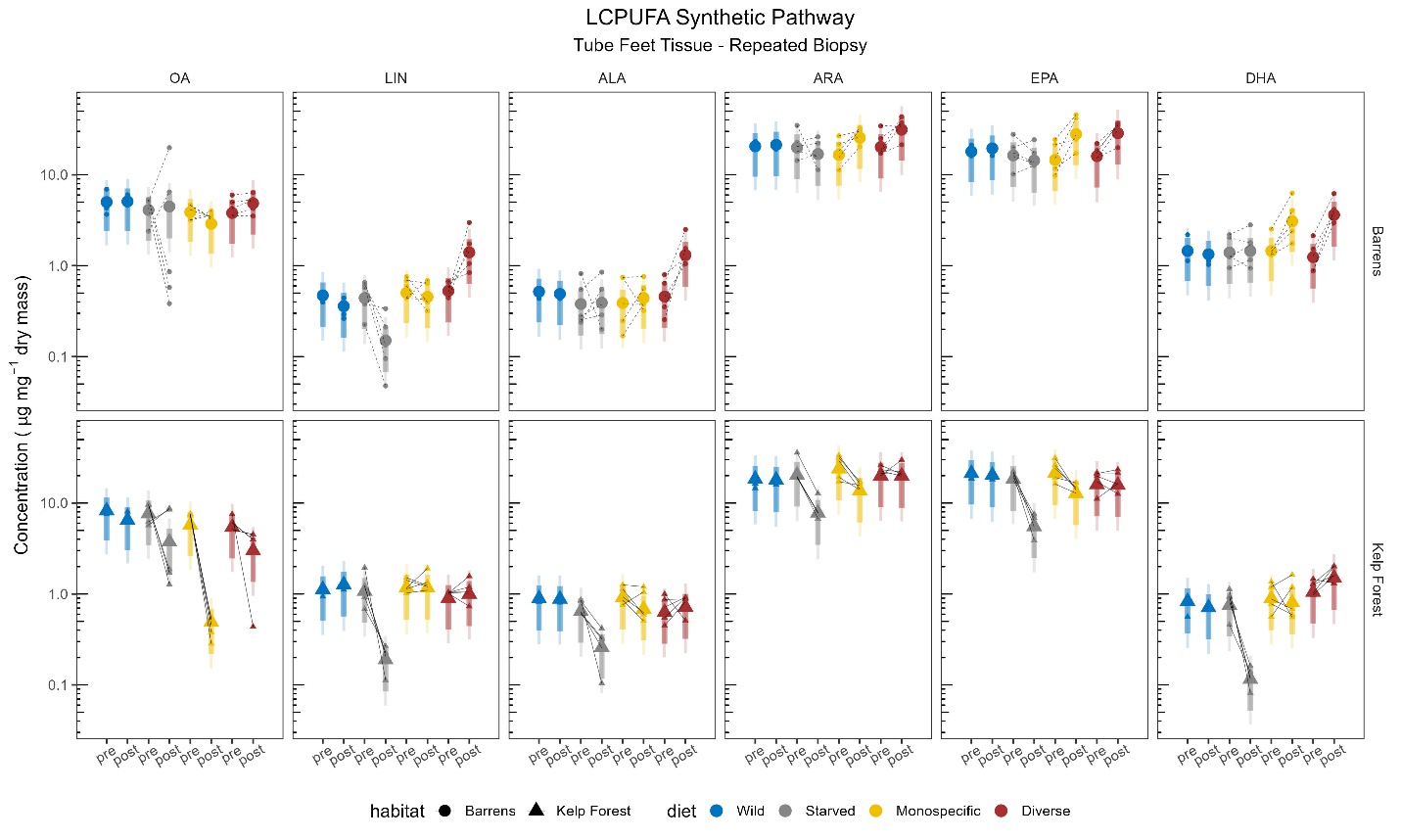

Figure S10: LCPUFA precursor assimilation longitudinal study – tube feet. Comparison of the concentration of six focal LCPUFA in repeated biopsies of tube feet tissue by treatment. The top row of panels represents data from barrens sea urchins, while the bottom row represents data from kelp forest urchins. Each column of panels represents data for a single focal fatty acid indicated by its abbreviation (left to right: OA, LIN, ALA, ARA, EPA, and DHA). Colors and solid shapes indicate diet treatments, where smaller faded symbols represent measured data points, larger solid symbols represent modeled mean values, and thin and thick vertical bars represent 95% and 80% credible intervals. Black lines connect pairs of data points within each diet to represent individual sea urchins subjected to repeated tube feet biopsies. The horizontal axis is delimited by two time points for each diet: “pre”, immediately prior to the experiment, and “post”, 33 days of treatment exposure.

Supplemental Tables

Table S1. Model selection for respiration analysis using leave one out cross validation stacking weights (2).

| Model | Predictors | LOO Stacking Weight |
| --- | --- | --- |
| **1** | **log(ash-free dry mass) * habitat * diet + (1\|chamber_id)** | **0.976** |
| 2 | log(ash-free dry mass) + habitat * diet + (1\|chamber_id) | 0.000 |
| 3 | log(ash-free dry mass) + habitat + diet + (1\|chamber_id) | 0.000 |
| 4 | log(ash-free dry mass) + habitat + (1\|chamber_id) | 0.024 |
| 5 | log(ash-free dry mass) + diet + (1\|chamber_id) | 0.000 |
| 6 | log(ash-free dry mass) + (1\|chamber_id) | 0.000 |

Table S2. Model selection for per capita feeding analysis based on algal biomass consumption using leave one out cross validation stacking weights (2).

| ID | Response units | Predictors | LOO Stacking Weight |
| --- | --- | --- | --- |
| 1 | algal biomass g dry urchin^-1^day^-1^ | log(test volume) * habitat * diet * feeding_date + (1\|individual_id) | 0.000 |
| 2 | algal biomass g dry urchin^-1^day^-1^ | log(test volume) + habitat * diet * feeding_date + (1\|individual_id) | 0.000 |
| 3 | algal biomass g dry urchin^-1^day^-1^ | log(test volume) + habitat + diet * feeding_date + (1\|individual_id) | 0.000 |
| 4 | algal biomass g dry urchin^-1^day^-1^ | log(test volume) + habitat * diet + feeding_date + (1\|individual_id) | 0.000 |
| 5 | algal biomass g dry urchin^-1^day^-1^ | log(test volume) + habitat + diet + feeding_date + (1\|individual_id) | 0.000 |
| 6 | algal biomass g dry urchin^-1^day^-1^ | log(test volume) + habitat + feeding_date + (1\|individual_id) | 0.125 |
| 7 | algal biomass g dry urchin^-1^day^-1^ | log(test volume) + diet + feeding_date + (1\|individual_id) | 0.000 |
| 8 | algal biomass g dry urchin^-1^day^-1^ | log(test volume) + feeding_date + (1\|individual_id) | 0.000 |
| **9** | **algal biomass g dry urchin^-1^day^-1^** | **log(spheroid_volume_ml) + diet * feeding_cycle_id + (1 \| individual_id)** | **0.875** |

Table S3. Model selection for per capita feeding analysis based on caloric consumption using leave one out cross validation stacking weights (2).

| ID | Response units | Predictors | LOO Stacking Weight |
| --- | --- | --- | --- |
| 1 | kcal urchin^-1^day^-1^ | log(test volume) * habitat * diet * feeding_date + (1\|individual_id) | 0.000 |
| 2 | kcal urchin^-1^day^-1^ | log(test volume) + habitat * diet * feeding_date + (1\|individual_id) | 0.000 |
| 3 | kcal urchin^-1^day^-1^ | log(test volume) + habitat + diet * feeding_date + (1\|individual_id) | 0.000 |
| 4 | kcal urchin^-1^day^-1^ | log(test volume) + habitat * diet + feeding_date + (1\|individual_id) | 0.000 |
| 5 | kcal urchin^-1^day^-1^ | log(test volume) + habitat + diet + feeding_date + (1\|individual_id) | 0.000 |
| 6 | kcal urchin^-1^day^-1^ | log(test volume) + habitat + feeding_date + (1\|individual_id) | 0.049 |
| 7 | kcal urchin^-1^day^-1^ | log(test volume) + diet + feeding_date + (1\|individual_id) | 0.000 |
| 8 | kcal urchin^-1^day^-1^ | log(test volume) + feeding_date + (1\|individual_id) | 0.000 |
| **9** | **kcal urchin^-1^day^-1^** | **log(spheroid_volume_ml) + diet * feeding_cycle_id + (1 \| individual_id)** | **0.951** |

Table S4. Model selection for per capita feeding analysis based on consumption of polyunsaturated fatty acids using leave one out cross validation stacking weights (2).

| ID | Response units | Predictors | LOO Stacking Weight |
| --- | --- | --- | --- |
| 1 | PUFA urchin^-1^day^-1^ | log(test volume) * habitat * diet * feeding_date + (1\|individual_id) | 0.000 |
| 2 | PUFA urchin^-1^day^-1^ | log(test volume) + habitat * diet * feeding_date + (1\|individual_id) | 0.000 |
| 3 | PUFA urchin^-1^day^-1^ | log(test volume) + habitat + diet * feeding_date + (1\|individual_id) | 0.000 |
| 4 | PUFA urchin^-1^day^-1^ | log(test volume) + habitat * diet + feeding_date + (1\|individual_id) | 0.000 |
| 5 | PUFA urchin^-1^day^-1^ | log(test volume) + habitat + diet + feeding_date + (1\|individual_id) | 0.120 |
| 6 | PUFA urchin^-1^day^-1^ | log(test volume) + habitat + feeding_date + (1\|individual_id) | 0.000 |
| 7 | PUFA urchin^-1^day^-1^ | log(test volume) + diet + feeding_date + (1\|individual_id) | 0.000 |
| 8 | PUFA urchin^-1^day^-1^ | log(test volume) + feeding_date + (1\|individual_id) | 0.000 |
| **9** | **PUFA urchin^-1^day^-1^** | **log(spheroid_volume_ml) + diet * feeding_cycle_id + (1 \| individual_id)** | **0.880** |

Table S5. Model selection for assimilation efficiency analysis using leave one out cross validation stacking weights (2).

| ID | Response units | Predictors | LOO Stacking Weight |
| --- | --- | --- | --- |
| 1 | Assimilation efficiency | log(test volume) * habitat * diet * feeding_date + (1\|individual_id) | 0.243 |
| 2 | Assimilation efficiency | log(test volume) + habitat * diet * feeding_date + (1\|individual_id) | 0.000 |
| 3 | Assimilation efficiency | log(test volume) + habitat + diet * feeding_date + (1\|individual_id) | 0.000 |
| 4 | Assimilation efficiency | log(test volume) + habitat * diet + feeding_date + (1\|individual_id) | 0.000 |
| **5** | **Assimilation efficiency** | **log(test volume) + habitat + diet + feeding_date + (1\|individual_id)** | **0.484** |
| 6 | Assimilation efficiency | log(test volume) + habitat + feeding_date + (1\|individual_id) | 0.062 |
| 7 | Assimilation efficiency ^-1^ | log(test volume) + diet + feeding_date + (1\|individual_id) | 0.000 |
| 8 | Assimilation efficiency ^-1^ | log(test volume) + feeding_date + (1\|individual_id) | 0.000 |
| 9 | Assimilation efficiency | log(spheroid_volume_ml) + diet * feeding_cycle_id + (1 \| individual_id) | 0.211 |

Table S6. Model selection for gonad mass analysis using leave one out cross validation stacking weights (2).

| Model | Predictors | LOO Stacking Weight |
| --- | --- | --- |
| 1 | log(test volume) * habitat * diet | 0.000 |
| 2 | log(test volume) + habitat * diet | 0.000 |
| **3** | **log(test volume) + habitat + diet** | **0.964** |
| 4 | log(test volume) + habitat | 0.000 |
| 5 | log(test volume) + diet | 0.000 |
| 6 | log(test volume) | 0.036 |

Table S7. Model selection for gonad lipid content analysis using leave one out cross validation stacking weights (2).

| Model | Predictors | LOO Stacking Weight |
| --- | --- | --- |
| **1** | **log(test volume) * habitat * diet** | **0.894** |
| 2 | log(test volume) + habitat * diet | 0.001 |
| 3 | log(test volume) + habitat + diet | 0.117 |
| 4 | log(test volume) + habitat | 0.000 |
| 5 | log(test volume) + diet | 0.106 |
| 6 | log(test volume) | 0.000 |

Table S8. Model selection for food conversion efficiency analysis using leave one out cross validation stacking weights (2).

| Model | Predictors | LOO Stacking Weight |
| --- | --- | --- |
| 1 | log(test volume) * habitat * diet | 0.000 |
| 2 | log(test volume) + habitat * diet | 0.000 |
| 3 | log(test volume) + habitat + diet | 0.000 |
| 4 | log(test volume) + habitat | 0.000 |
| **5** | **log(test volume) + diet** | **1.000** |
| 6 | log(test volume) | 0.000 |

Table S9. Model selection for kelp biomarker analysis using leave one out cross validation stacking weights (2).

| Model | Predictors | LOO Stacking Weight |
| --- | --- | --- |
| 1 | log(test volume) * habitat * diet | 0.092 |
| 2 | log(test volume) + habitat * diet | 0.092 |
| **3** | **log(test volume) + habitat + diet** | **0.760** |
| 4 | log(test volume) + habitat | 0.000 |
| 5 | log(test volume) + diet | 0.057 |
| 6 | log(test volume) | 0.000 |

Table S10. Model selection for biofilm biomarker analysis using leave one out cross validation stacking weights (2).

| Model | Predictors | LOO Stacking Weight |
| --- | --- | --- |
| 1 | log(test volume) * habitat * diet | 0.001 |
| **2** | **log(test volume) + habitat * diet** | **0.965** |
| 3 | log(test volume) + habitat + diet | 0.000 |
| 4 | log(test volume) + habitat | 0.000 |
| 5 | log(test volume) + diet | 0.000 |
| 6 | log(test volume) | 0.034 |

Table S11. Comparison of nutritional quality among macroalgae in controlled diets. Values measured in the current study include the columns for fatty acid content expressed as mean concentration (μg g^-1^ dy wt) with parenthetical 95% credible intervals (calculated using the R package brms(3)). The rest of the nutritional parameters are derived from the literature.

| **Species** | **ω-3 (μg g^-1^ dry wt)** | **ω-6 (μg g^-1^ dry wt)** | **DHA-Precursors (μg g^-1^ dry wt)** | **(kcal g^-1^ dry wt)** | **C:N** | **protein** |
| --- | --- | --- | --- | --- | --- | --- |
| *Chondracanthus corymbiferus* | 8.53 (5.47-13.50) | 14.06 (8.81-23.01) | 28.21 (18.38-44.16) | 3.34^a^ (CV 0.9) | (Rhodophyta) 10.2 (SE 0.5)^d^ | (*Chondrus crispus*) 11-18^f^ |
| *Dilsea californica* | 6.89 (4.49-11.57) | 14.51 (9.34-24.39) | 22.46 (15.20-36.23) | (*Opuntiella californica*) 3.53^a^ | (Rhodophyta) 10.2 (SE 0.5)^d^ | (*Chondrus crispus*) 11-18^f^ |
| *Nereocystis luetkeana* | 1.99 (1.26-3.46) | 5.57 (3.61-9.67) | 11.10 (7.43-18.53) | 4.00^b^ (SD 1.1) | 10^d^ | (*Laminaria saccharina*) 6-11^f^ |
| *Ulva spp.* | 7.12 (4.56-12.06) | 7.01 (4.51-12.13) | 16.60 (11.01-26.81) | 3.03^c^ (SE 0.08) | (Chlorophyta) 9.4 (SE 1.4)^d^ | 15-25^f^ |
|  | ^a^Paine and Vadas (1969)(4), ^b^Dethier et al (2014)(5), ^c^Lamare and Wing (2001)(6), ^d^Peters *et al* (2005)(7), ^e^Rosell and Srivastava (1985)(8), ^f^Morrissey, Kraan, and Guiry (2001)(9) | | | | | |

Inverse problem: modeling gonad mass and feeding rates jointly

3. Gonad mass model:

Equation S1

$$g_{i}^{*}= \beta_{0}^{g}+ \beta_{1}^{g}\log{body size}_{i}+\beta_{2}^{g}H_{i}+\beta_{3}^{g}D_{i}+\beta_{4}^{g}\left( H_{i}\times D_{i} \right)+\beta_{5}^{g}f_{i}^{*}+\epsilon_{i}^{g}$$

Where:

- $g_{i}^{*}$ is the true (latent) gonad mass for the $i$th individual.
- $\log({body size}_{i})$ is the log-transformed body size for the $i$th individual.
- $H_{i}$ and $D_{i}$ are the categorical variables for habitat and diet, respectively.
- $f_{i}^{*}$ is the true (latent) feeding rate for the $i$th individual.
- $\epsilon_{i}^{g}$ is the residual error term for the gonad mass model.
- $\beta_{0}^{g}$ is the intercept for the gonad mass model, and $\beta_{1}^{g},\beta_{2}^{g},\beta_{3}^{g},\beta_{4}^{g},\beta_{5}^{g}$ are the regression coefficients for the corresponding predictors.

The observed gonad mass $g_{i}$ is related to the latent gonad mass $g_{i}^{*}$ by:

Equation S2

$$g_{i}= g_{i}^{*}+{measurement error}_{i}^{g}$$

4. Feeding rate model:

Equation S3

$$f_{i}^{*}= \beta_{0}^{f}+ \beta_{1}^{f}\log{body size}_{i}+\beta_{2}^{f}H_{i}+\beta_{3}^{f}D_{i}+\beta_{4}^{f}\left( H_{i}\times D_{i} \right)+\epsilon_{i}^{f}$$

- $f_{i}^{*}$ is the true (latent) feeding rate for the $i$th individual.
- $\log({body size}_{i})$ is the log-transformed body size for the $i$th individual.
- $H_{i}$ and $D_{i}$ are the categorical variables for habitat and diet, respectively.
- $f_{i}^{*}$ is the true (latent) feeding rate for the $i$th individual.
- $\epsilon_{i}^{f}$ is the residual error term for the feeding rate model.
- $\beta_{0}^{f}$ is the intercept for the feeding rate model, and $\beta_{1}^{f},\beta_{2}^{f},\beta_{3}^{f},\beta_{4}^{f},\beta_{5}^{f}$ are the regression coefficients for the corresponding predictors.
- The observed gonad mass $g_{i}$ is related to the latent gonad mass $g_{i}^{*}$ by:

Equation S4

$$f_{i}= f_{i}^{*}+{measurement error}_{i}^{f}$$

Trophic modification and assimilation of fatty acids in *M. franciscanus*

Our experimental results combined with data from the wild source populations suggest that *M. franciscanus* retains in gonadal tissues several precursor fatty acids involved in the biosynthesis of DHA directly from ingested macroalgae**.** ω3 fatty acids like DHA and its precursor molecules ALA and EPA are known to affect herbivore fitness via reproductive physiology and survival (reviewed in(12)). The four species of macroalgae offered in the experiment each represented a separate family and had distinct fatty acid compositions, consistent with previous findings for temperate macrophytic algae(13), and had very low concentrations of DHA, but the gonadal tissues of experimental subjects contained DHA at concentrations comparable to levels of its biosynthetic precursors (**figure** **S8**). Legacy effects of DHA-rich food availability in kelp forest urchins could explain the discrepancy between low DHA availability in the experimental diets and high DHA and DHA-precursor concentrations in *M. franciscanus* tissues. However, barrens urchins began the experiment with low DHA concentrations in their gonadal tissues, then had higher concentrations at the end of the experiment when fed macroalgae rich in DHA precursors, suggesting a capacity to synthesize DHA from precursors via multiple enzymatic steps involving front end desaturases and PUFA elongases (see(12) for synthesis of fatty acid biosynthetic pathways). Barrens urchins that were experimentally starved showed no change in DHA, indicating they did not synthesize DHA *de novo*. Moreover, the concentration of EPA in urchin tissues was exceptionally high relative to the other DHA precursors ARA and ALA even though ARA and ALA occurred in comparable concentrations to EPA in the experimental macroalgal diets. This relative difference provides support both for the retention of EPA from consumption of macroalgae and for the presence of a conserved biosynthetic pathway wherein DHA can be converted to the shorter EPA via beta oxidation, as has been shown in aquatic invertebrates like copepods and ragworms (14-16). As has been demonstrated in experimentally starved copepods (11), kelp urchins appeared to mobilize multiple PUFAs when starved, whereas barrens urchins only mobilized LIN, a PUFA found in high concentrations in *Ulva sp.* These declines in specific PUFAs, particularly LIN in barrens urchins, concomitant with starvation-related reductions in overall lipid content, indicate the possibility that LIN may be limiting for the nutritional needs of *M. franciscanus*.

Our longitudinal study of repeated tube feet biopsies resulted in qualitatively similar treatment specific shifts in fatty acid profiles relative to gonadal tissues with respect to OA, LIN, and ALA, though tube feet had at least three orders of magnitude lower lipid content (**figure S9**). Concentrations of LIN, ALA, ARA, EPA, and DHA all decreased in final tube feet biopsies in urchins from the kelp forest that were experimentally starved. Kelp forest urchins fed a diverse diet exhibited an increase in the concentration of DHA in final tube feet biopsies relative to those fed a monospecific diet or those from the wild population. Aside from DHA, final tube feet biopsies from kelp forest urchins fed either a diverse or monospecific diet did not change substantially with respect to focal fatty acids relative to their initial values. Barrens urchins that were experimentally starved showed a decrease in the concentration of LIN in final tube feet biopsies relative to their initial values but showed almost no change in other focal FAs. Barrens urchins that were fed a diverse diet exhibited increases in concentrations of LIN, ALA, ARA, EPA, and DHA in final tube feet biopsies while those fed a monospecific diet showed increases in concentrations of ARA, EPA, and DHA, but not LIN or ARA. In general, the fatty acid composition of urchin tissues reflected a clear starvation signal related to bacterially derived fatty acids, whereas algal fatty acid diet tracers were more nuanced. A focal fatty acid summation model for repeated tube feet biopsies including a family-level effect of individual plus population-level effects of body size and the three-way interaction between habitat, diet, and focal fatty acid was decisively favored over simpler model structures (PP = 1.000, BF >100, SE = 1%)
